# Genetically encoded control of *in vitro* transcription-translation coupled DNA replication

**DOI:** 10.1101/2025.07.08.663768

**Authors:** Sebastian Barthel, Maximilian Hoffmann-Becking, Islomjon G Karimov, Tobias J Erb

**Affiliations:** Department of Biochemistry & Synthetic Metabolism, Max Planck Institute for Terrestrial Microbiology, Karl-von-Frisch-Str. 10, 35043 Marburg, Germany; Center for Synthetic Microbiology (SYNMIKRO), Philipps University Marburg, Karl-von-Frisch Str. 14, 35043 Marburg, Germany

**Keywords:** transcription-translation coupled DNA replication (TTcDR), PURE system, genetically encoded system control, genetic circuit, *in vitro* systems, synthetic cell

## Abstract

The bottom-up reconstruction of cellular functions has gained increasing attention for studying biological complexity, and for developing advanced biotechnological tools, including synthetic cells. A fundamental challenge is the ability to control and replicate DNA-encoded information within basic *in vitro* transcription-translation (IVTT) systems. Here, we constructed a transcription-translation coupled DNA replication (TTcDR) system that is based on a modified PURE (<u>P</u>rotein synthesis <u>U</u>sing <u>R</u>ecombinant <u>E</u>lements) IVTT system and Φ29 DNA polymerase, which is controlled by external signals. To this end, we first established and characterized a PUREfrex 1.0-based TTcDR system. We then constructed and optimized TetR-based control of TTcDR activity, either by DNA-encoding TetR or by supplying purified TetR. Our final DNA-encoded TetR circuit allows ∼1000-fold DNA replication, ∼100-fold repression, and ∼4-fold induction with anhydrotetracycline. Our results demonstrate the potential and challenges of controlling *in vitro* DNA replication, for example for the evolution of *in vitro* systems.

**Graphical Abstract:** 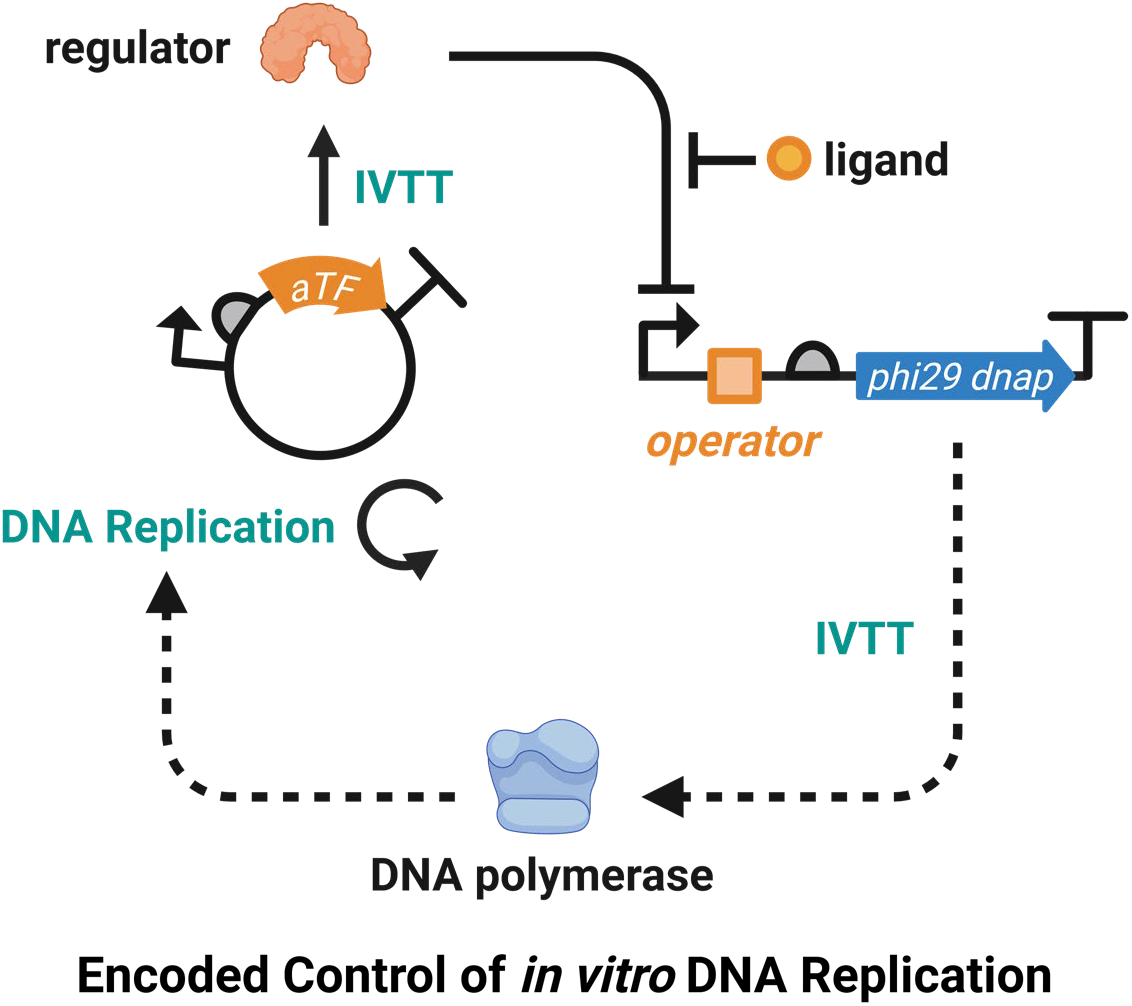

## Introduction

The bottom-up reconstruction of the fundamental functions of living cells is a central goal of synthetic biology. These efforts aim at unraveling the underlying operating principles of living systems, deepen our understanding of biological complexity, and lay the foundation for the development of advanced bio(techno)logical systems, such as artificial organelles and synthetic cells (1–3). One important tool in these efforts is the <u>P</u>rotein synthesis <u>U</u>sing <u>R</u>ecombinant <u>E</u>lements (PURE) system (4). This *in vitro* transcription-translation system, reconstituted from *Escherichia coli* proteins, ribosomes and tRNAs, provides researchers with the opportunity to operate processes of the central dogma outside of the context of living cells.

A fundamental process of living cells is their ability to replicate DNA-encoded information, which enables proliferation and evolution (1, 5–7). Recently, several *in vitro* DNA replication schemes have been reported that operate in the PURE system and are generally referred to as transcription-translation coupled DNA replication (TTcDR). All these approaches utilize Φ29 DNA polymerase. In a stand-alone fashion, this enzyme either catalyzes rolling circle amplification (RCA) of circular DNA to produce concatemeric DNA (8–13), or is capable of replicating linear DNA, when assisted by auxiliary proteins of the Φ29 replication machinery (5, 6, 14, 15). The Ichihashi and Mutschler groups have recently established stand-alone Φ29 DNA polymerase-based TTcDR reactions within modified PURE systems (8–10).

The precise control of genetic networks in response to external signals is another fundamental process of living systems. While sophisticated genetic circuits are widely used *in vivo* (16–19) and increasingly in cell extract-based transcription-translation systems (20–23), few genetic circuits have been reported in the PURE system, so far. These *in vitro* circuits are typically based on a variety of nucleic acid-based regulatory elements that respond to small molecules (24, 25), temperature (26–28), or light (29–31). Allosteric transcription factors (aTFs), which are commonly used in the design of genetic circuits *in vivo*, have only been used rarely *in vitro* (32, 33), even though they cover a wider range of ligands (34) and offer increased dynamic ranges compared to riboswitches (35–37) or optogenetics-based systems (38, 39).

In this study, we sought to construct a controllable TTcDR system that is based on a genetically encoded circuit. To that end, we first tested different Φ29 DNA polymerase stand-alone TTcDR systems for their DNA replication, *in vitro* transcription and *in vitro* transcription-translation capacities. We further established a tetracycline repressor (TetR)-based genetic circuit with fluorescent reporter and DNA replication readout under TTcDR conditions. Finally, we tested and engineered different Φ29 DNA polymerase variants to further increase TTcDR activity and improve genetic circuit performance. Overall, our work provides a controlled *in vitro* DNA replication system that responds to external signals, opening the possibility for integration of more complex functions, such as metabolic networks (40–42) or Darwinian evolution into *in vitro* transcription-translation systems (3, 6, 43), which will be a critical step toward constructing more complex systems, such as synthetic cells in the future (1, 2, 44).

## Results

### Characterizing and optimizing a transcription-translation coupled DNA replication (TTcDR) system

To establish a TTcDR system, we sought to characterize and optimize the commercially available PUREfrex 1.0 system and different commercial and customized energy mixes (9, 10) for DNA replication, *in vitro* transcription (IVT), and *in vitro* transcription-translation (IVTT) capacities (Figure 1).

**Figure 1:**
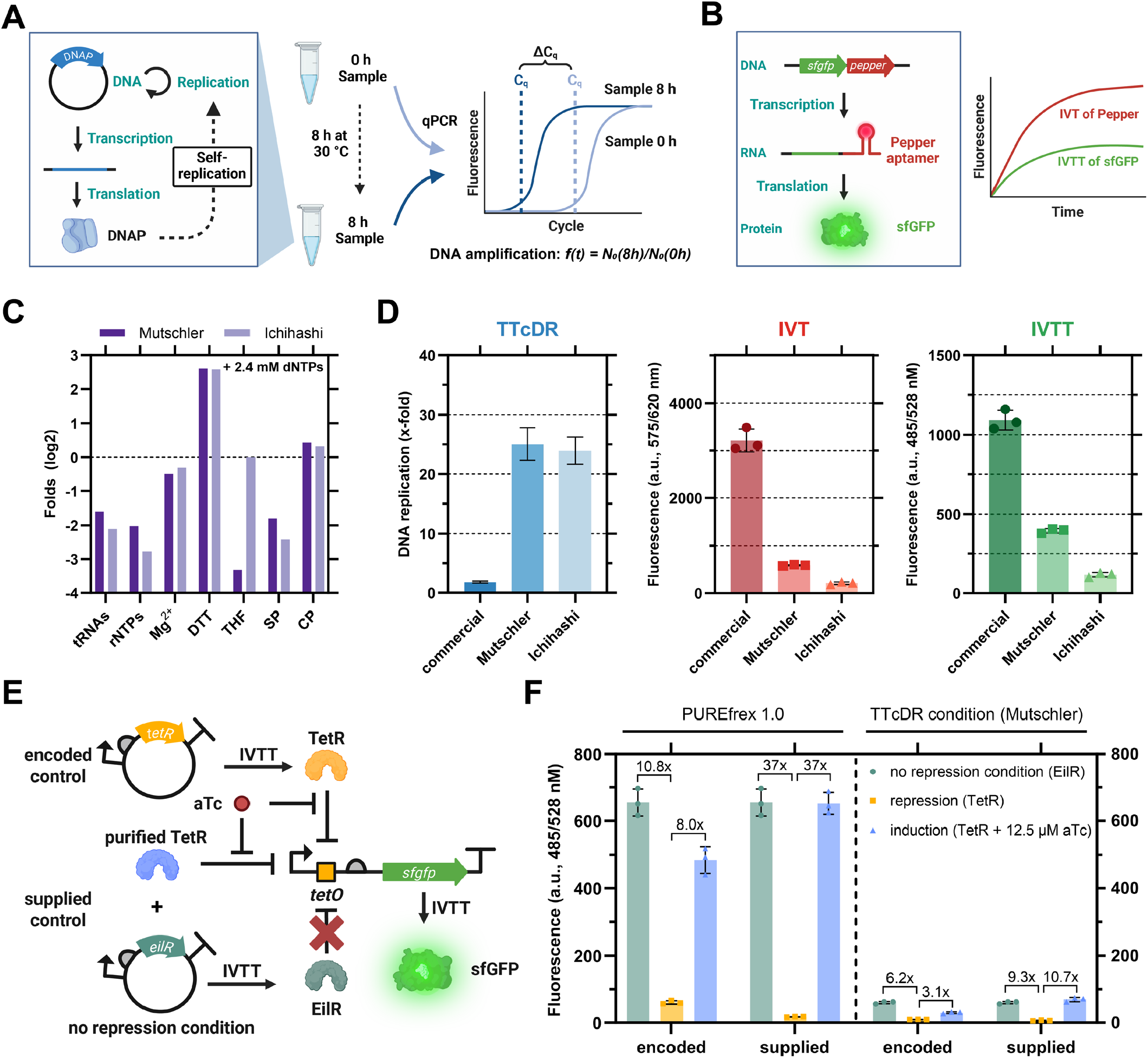
Characterization of a stand-alone Φ29 DNA polymerase-based transcription-translation-coupled DNA replication (TTcDR) system based on the commercial PUREfrex 1.0 system and characterization of a TetR-based genetic circuit with IVT(T) output. **A:** Schematic of TTcDR experiments based on Φ29 DNA polymerase-driven self-replication and qPCR analysis. Φ29 DNA polymerase is produced in the IVTT system from plasmid, which is in turn self-replicated by Φ29 DNA polymerase. DNA replication is analyzed by comparing DNA quantities measured by qPCR of samples taken before and after incubation at 30°C. **B:** Schematic of IVTT experiments based on a two-color fluorescence plate reader assay using the Pepper aptamer with HBC620 ligand to measure *in vitro* transcription (IVT) in the red channel, and sfGFP to measure *in vitro* transcription-translation (IVTT) in the green channel. **C:** Major differences in the composition of the two custom EMs (9, 10) compared to the PUREfrex 1.0 energy mix (48). See Figure S1E for a complete comparison. SP: spermidine. CP: creatine phosphate. **D:** Comparison of TTcDR activity (not baseline corrected), and IVT(T) capacities of PUREfrex 1.0-based TTcDR systems with either commercial or customized energy mixes. More detailed data are shown in Figure S1-3. **E:** Schematic of the genetic circuit. TetR is produced from plasmid DNA to repress the PT7-*tetO* promoter on the linear DNA template, which controls Pepper aptamer expression and sfGFP production. In the presence of aTc, TetR unbinds PT7-*tetO* and the circuit is induced. For encoded control, either TetR (repressing) or EilR (non-repressing) were co-produced. For supplied control, 500 nM purified TetR was added to EilR co-producing samples. **F:** IVTT output from the TetR-based genetic circuit in PUREfrex 1.0 and under standard TTcDR conditions. Fold changes of repression and induction are shown above the bars. Full IVT data and IVTT kinetics are shown in Figure S5. Data are the mean of n = 3 replicates ± SD.

To test for DNA replication activity, we cloned the Φ29 bacteriophage DNA polymerase gene *p2* under the control of a T7 phage promoter and produced Φ29 DNA polymerase in the respective TTcDR system. After production, Φ29 DNA polymerase begins to self-replicate its plasmid by rolling circle DNA amplification (RCA). We measured DNA replication by quantitative polymerase chain reaction (qPCR) (Figure 1A) and observed poor DNA amplification in the TTcDR systems with the commercial energy mix (Figure 1D, S1-2). This observation is consistent with previous reports suggesting inhibition of DNA replication, especially by high concentrations of transfer ribonucleic acid (tRNA) and ribonucleoside triphosphates (rNTP) (8, 9). In contrast, we observed 25-fold and 24-fold DNA amplification with customized energy mixes of the Mutschler and Ichihashi labs that are reduced in tRNA and rNTPs. When testing for nucleic acid stability, we observed some basic nuclease activity at the lower limit of detection, indicating negligible degradation of RNA and DNA in our *in vitro* system (Figure S1F).

Next, we investigated the IVTT capacity of the PURE systems with the commercial and TTcDR-compatible energy mixes. For this, we used a two-color fluorescent plate reader assay based on a linear DNA template encoding the Pepper RNA aptamer to quantify transcriptional activity and superfolder green fluorescent protein (sfGFP) to monitor transcriptional-translational activity (Figure 1B) (45, 46). Compared to the commercial energy mix, we observed 5-15 fold reduced IVT output, and 3-9-fold reduced IVTT output with the Mutschler and Ichihashi energy mixes, respectively (Figure 1D, S3). Yet, these loses in IVT and IVTT output were still compensated by improved TTcDR activity of the customized energy mixes (see above).

In the following, we tested how the concentration of PUREfrex 1.0 proteins and ribosomes affected IVT and IVTT capacities of the different energy mixes (10). When increasing protein and ribosome concentrations by 1.5-fold and 2-fold, respectively, we observed improved IVT and IVTT kinetics for all energy mixes tested (Figure S4A,B). However, under these conditions DNA replication was decreased between 2-8-fold in the customized energy mix (Figure S4C), reminding of similar observations when trying to balance between translation and RNA replication (47).

Finally, we also tested the performance of the PUREfrex 2.0 and PURExpress-based TTcDR systems with the three energy mixes. Both PURE systems showed similar behavior as PUREfrex 1.0 (Figure S1), however, they are 1.5 - 3x more expensive and less well described in their composition. Thus, we decided to continue with the normal concentrations of proteins and ribosomes in PUREfrex 1.0 using the Mutschler energy mix, in the following referred to “standard TTcDR condition”. This condition has the advantage that the PUREfrex 1.0 composition is fully known (48), which provides the possibility for additional optimizations of individual components, if required.

### Characterization of a TetR-based genetic circuit on transcriptional and translational level

Next, we constructed a genetic circuit with fluorescence readout for IVT and IVTT in our TTcDR system. To that end, we chose TetR from *Escherichia coli*, which was previously used to construct a genetic circuit and a genetic oscillator in the PURE system (32, 33). In our genetic circuit, we co-expressed *tetR* from a plasmid (pTetR) and the *sfGFP-pepper* reporter under the control of a T7-*tetO* promoter from a linear DNA template (Figure 1E). Upon production, TetR represses transcription of the T7-*tetO* promoter unless anhydrotetracycline (aTc), a ligand of TetR, is present that induces transcription through de-repression (49). As a non-regulated expression control, we chose EilR from *Enterobacter lignolyticus*, a TetR family transcriptional repressor of similar molecular weight as TetR (pEilR) that does not bind *tetO* (50). When expressing our genetic circuit at equimolar concentrations of DNA under PUREfrex 1.0 conditions, we observed an 11-fold repression of sfGFP production in samples co-producing TetR compared to control samples co-producing EilR (Figure 1F, S5).

Note that genetically encoded circuits in cell-free systems show some leakage due to the absence of transcriptional repressors in the initial phase of PURE reactions (i.e., before the repressor is produced in sufficient amounts). To elucidate the effect of the temporal delay on repression, we supplied 500 nM purified TetR and observed a 37-fold repression of sfGFP production, which is a 3.5-fold improvement compared to TetR co-producing circuits. Next, we tested induction of the system. When adding 12.5 µM aTc to the genetic circuit, we observed 8-fold increased sfGFP signal output when co-producing TetR and 37-fold increased sfGFP signal when supplying 500 nM purified TetR.

When testing the gene circuit under “standard TTcDR conditions”, we observed a significantly lower dynamic range, compared to PUREfrex 1.0 conditions, i.e., 6-fold repression and 3-fold induction of sfGFP signal when co-producing TetR, and 9-fold repression and 11-fold induction with 500 nM purified TetR (Figure 1F, S5). We hypothesized that under TTcDR conditions, gene expression from plasmid DNA templates is preferred over linear DNA templates, causing a decrease in reporter production from the linear template. This was further corroborated by the fact that in PUREfrex 1.0, IVT and IVTT output from a linear DNA template dropped by ∼2x when adding an equimolar concentration of plasmid DNA, but 4-6x under TTcDR conditions (Figure S5F). Further experiments confirmed a preference for circular DNA templates under standard TTcDR conditions but not in PUREfrex 1.0 (Note S1, Figure S6).

### Encoded control of DNA replication by genetic circuitry

Next, we coupled the TetR-based genetic circuit to DNA replication. To that end, we placed the Φ29 DNA polymerase gene *p2* under the control of a T7-*tetO* promoter on a linear DNA template and co-produced either TetR or EilR from plasmid DNA (Figure 2A). We next titrated the linear-to-circular DNA ratio in TetR co-producing TTcDR reactions for best repression. With increasing linear-to-circular DNA ratio, DNA replication and repression of the genetic circuit increased to ∼60-fold DNA replication and 20-fold repression (Figure 2B). Next, we tested the induction of TetR-based genetic circuit with 12.5 µM aTc and a 16-fold linear-to-circular DNA ratio. We observed ∼16-fold repression and ∼2-fold induction, when encoding TetR, and ∼50-fold repression and ∼30-fold induction, when supplying 500 nM purified TetR (Figure 2C). When we increased the aTc concentration to 50 µM, we observed a decrease in DNA replication, indicating inhibition of the system (Figure S7).

**Figure 2:**
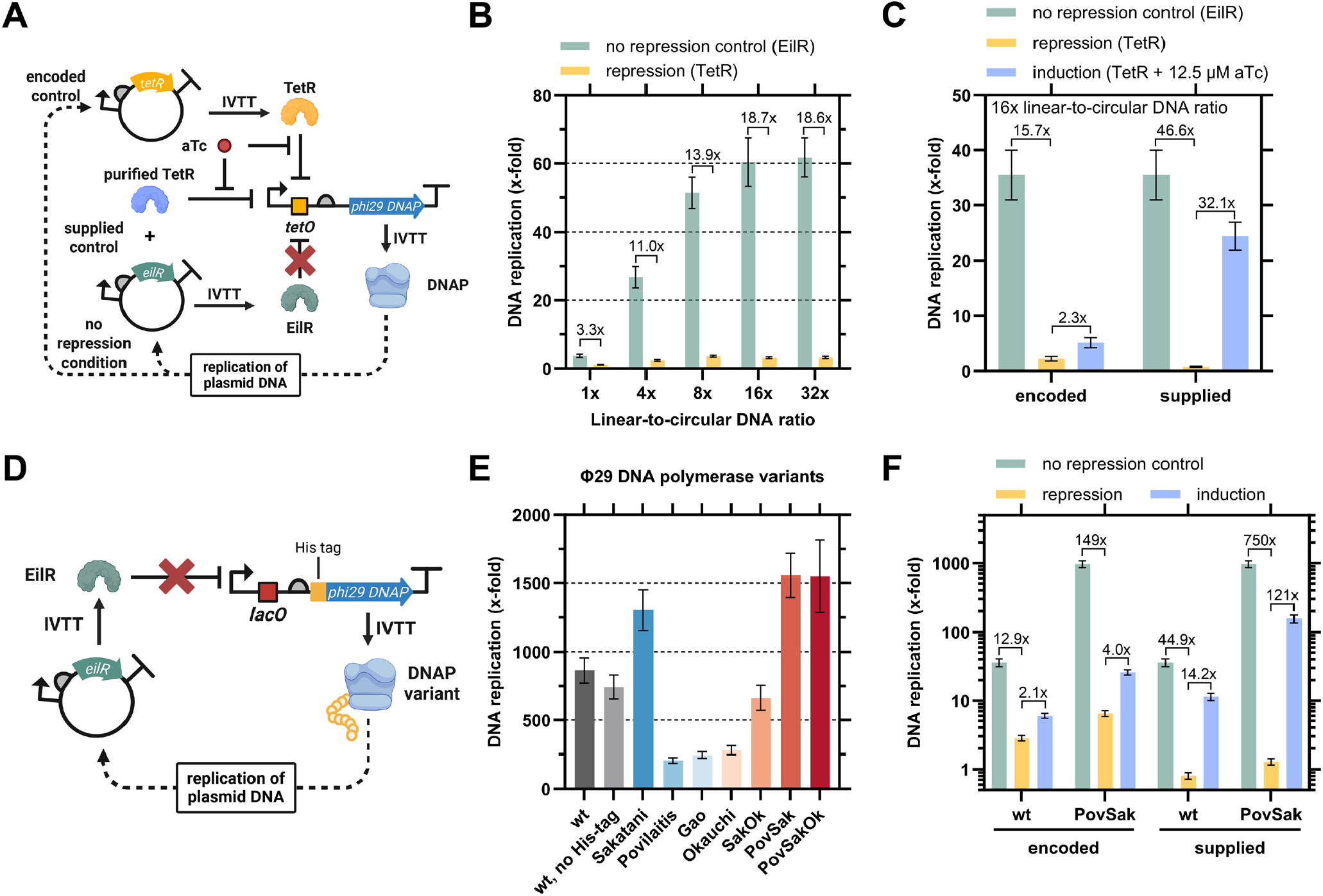
Characterization of a TetR-based genetic circuit with TTcDR output, and testing of Φ29 DNA polymerase mutants to improve circuit performance. **A:** Schematic of the genetic circuit. TetR is produced from plasmid DNA to repress the PT7-*tetO* promoter on the linear DNA template, which controls the production of Φ29 DNA polymerase. In the presence of aTc, TetR unbinds PT7-*tetO* and the circuit is induced, leading to the production of DNA polymerase that replicates the plasmid DNA. For encoded control, either TetR (repressing) or EilR (non-repressing) were co-produced. For supplied control, 500 nM purified TetR was added to samples co-producing EilR. **B:** TTcDR output from the genetic circuit at various ratios of linear DNA (fixed at 4 nM) to circular DNA (0.125 nM – 4 nM) co-producing either EilR (non-repressing) or TetR (repressing). **C:** TTcDR output under encoded and supplied control. **D:** Schematic of testing Φ29 DNA polymerase mutants. The Φ29 DNA polymerase is produced from linear DNA under control of a PT7-*lacO* promoter to replicate plasmid DNA encoding EilR. **E:** TTcDR output of Φ29 DNA polymerase mutants. Mutant names correspond to the first author of their respective publication. All mutants are N-terminally His-tagged, unless otherwise noted. **F:** Comparison of Φ29 DNA polymerase wild-type and the His-tagged PovSak mutant in TetR-controlled TTcDR. Fold changes of repression and induction are shown above the bars. **C-F:** All samples used 16x linear-to-circular DNA ratios. Data are the mean of n = 3 replicates ± SD. TTcDR data are baseline corrected with data from inactive Φ29 DNA polymerase samples (56).

### Φ29 DNA polymerase mutant improves genetic circuit-controlled TTcDR

When encoding a weaker RBS for Φ29 DNA polymerase translation, we observed reduced TTcDR performance (Figure S8). Thus, we suspected that increasing Φ29 DNA polymerase activity would improve circuit performance. We therefore tested four Φ29 DNA polymerase variants described in literature (51– 54) and three combinations of mutations (Table S7). All seven variants were cloned under control of a P_T7_-*lacO* promoter with an N-terminal His-tag to ensure the same N-terminal sequence, as the first amino acids of open reading frames were reported to significantly affect protein synthesis *in vitro* (55). When testing the seven variants for replication of the EilR-encoding plasmid (Figure 2D), we observed improved DNA replication for three mutants compared to wild-type Φ29 DNA polymerase, with the combined mutant “PovSak” performed best (∼2-fold over wild-type) (Figure 2E).

Notably, we also observed that overall DNA replication in our screen was significantly higher than in previous experiments. We hypothesized that an increase in promoter activity of PT7-*lacO* over PT7-*tetO* and an increased translation rate due to a favorable N-terminal sequence could cause the improvement, and indeed observed additive 10-fold improvements by the PT7-*lacO* promoter and N-terminal His-tag sequence (Figure S9).

Finally, we compared the His-tagged PovSak mutant with wild-type Φ29 DNA polymerase in the TetR circuit side-by-side. Strikingly, the PovSak mutant showed ∼970-fold total DNA replication and only 6.5-fold total DNA replication when co-producing TetR (Figure 2F). This results in an improved repression of 149-fold with the His-tagged PovSak mutant versus 13-fold with the wild-type DNA polymerase. In addition, induction with 12.5 µM aTc improved DNA replication from 2.1-fold to 4-fold, indicating that the improved Φ29 DNA polymerase activity overall improved the performance of the circuit. Alternatively, the TetR-supplemented system provides 750-fold repression and ∼120-fold induction for applications in which supplied control of TTcDR is suitable. Overall, the His-tagged PovSak mutant drastically improved the genetically encoded control of the TTcDR system to provide high DNA replication and high repression.

## Discussion

Here, we report the first control of transcription-translation coupled DNA replication (TTcDR) using genetic circuitry. Our system makes use of the aTF TetR from *E. coli*, which had been prototyped in IVT(T) systems (32, 33, 49) and which we successfully coupled to control DNA replication. Our encoded system is capable of ∼1000-fold DNA replication under non-repressing conditions, shows ∼150-fold repression under TetR co-producing conditions, and a 4-fold response to the inducer anhydrotetracycline. If TetR is supplemented from the beginning, the system becomes even more controllable with 750-fold repression and a 120-fold response to aTc.

All of our experiments showed a significant difference in repression and induction between co-produced TetR and 500 nM purified TetR. We attribute the reduced repression in co-production circuits to the absence of aTF in the initial phase of transcription-translation of the reporter protein. Concerning the circuit induction, the co-producing system shows a 4-fold response to 12.5 µM aTc, which clearly leaves room for improvement to use such a system in synthetic cells or other applications. We argue that this is a two-sided problem. On the one hand, aTF production is not negatively regulated and thus continues until the PURE system is depleted. After poor repression in the initial phase of the reaction, repression becomes very strong, requiring more ligand to completely induce the system. On the other hand, reconstituted *in vitro* systems are prone to crosstalk with or inhibition by additional components such as aTc (Figure S7) (57) and other ligands. For example, we have previously mapped the effects of 13 non-enzyme components of a synthetic metabolic system that drastically inhibited *in vitro* transcription and the PURE system (42, 58).

The control of DNA replication within the PURE system using an encoded genetic circuit provides a new opportunity to integrate metabolic *in vitro* modules in the PURE system via Darwinian evolution, such as synthetic metabolism (40–42) or lipid biogenesis (5, 43). This requires coupling the activity of the metabolic *in vitro* module to DNA replication (3). Our proof-of-concept study showed that genetic circuits based on aTFs could be suitable to transduce metabolic activity to DNA replication.

## Materials and Methods

### Reagents

Unless otherwise noted, chemicals were purchased from Merck KGaA (Darmstadt, Germany) and Carl Roth GmbH (Karlsruhe, Germany). Commercial enzymes and bioreagents were purchased from New England Biolabs (Frankfurt am Main, Germany). Commercial PURE systems were purchased from GeneFrontier (PUREfrex 1.0, PUREfrex 2.0; Kashiwa, Japan) and New England Biolabs (PURExpress). Nuclease detection kits were purchased from Jena Biosciences (Jena, Germany).

### Strains and growth media

For molecular cloning, *Escherichia coli* NEB 5a was grown in lysogeny broth (LB) supplemented with an appropriate antibiotic (100 µg/ml ampicillin or 34 µg/ml chloramphenicol). For protein production of TetR, *E. coli* BL21 (DE3) was grown in terrific broth (TB) supplemented with 50 µg/ml ampicillin. All strains used are listed in Table S1.

### Plasmid assembly and preparation of linear DNA templates

Oligonucleotides were purchased from Merck KGaA. Synthetic dsDNA was purchased from Twist Bioscience (South San Francisco, CA, USA). Sanger sequencing was performed by MicroSynth (Göttingen, Germany).

Plasmids were generated by Golden Gate Assembly (GGA). Level 0 and level 1 plasmids were assembled using the modular cloning system proposed by Stukenberg et al. (59). Plasmids encoding intramolecular circuits were assembled from PCR-amplified fragments with GGA overhangs and an isolating spacer sequence between the two transcriptional units, using the respective level 1 plasmids encoding the aTF gene as a backbone. A mixture of 0.5 nM vector DNA and 2 nM of each insert DNA was assembled using 0.5 U/µl Esp3I (level 0) or 1 U/µl BsaI-HFv2 (level 1 & intramolecular circuits) and 40 U/µl T4 ligase in 1× T4 ligase buffer. GGA reactions were cycled 15 times for 1.5 min at 37°C and 3 min at 16°C. Enzymes were heat-inactivated for 5 min at 50°C and 10 min at 80°C. The GGA product was transformed into chemically competent *E. coli* NEB 5-alpha cells, and individual clones were verified by Sanger sequencing. Plasmids were purified using the NucleoSpin Plasmid kit (Macherey-Nagel, Düren, Germany), according to the manufacturer’s instructions and checked for circularity on an agarose gel.

All linear DNA templates were prepared by polymerase chain reaction (PCR) amplification from the respective plasmids using Q5 DNA polymerase, according to the manufacturer’s instructions. All amplified DNA fragments were purified using the NucleoSpin Gel and PCR Clean-up kit (Macherey-Nagel), according to the manufacturer’s instructions. All DNA concentrations were calculated from absorbance measurements at 260 nm (A260) using a NanoDrop2000 spectrophotometer (Thermo Scientific, Waltham, MA, USA). All plasmids and linear DNA templates, as well as oligonucleotides used to generate linear DNA templates, are listed in Tables S2, S3 and S4, respectively. DNA sequences are provided on FigShare (Data Availability).

### Production and purification of TetR

TetR was produced in an *E. coli* BL21 (DE3) strain harboring plasmid pJBL701 (49). First, a preculture was inoculated in TB, supplemented with 50 µg/mL kanamycin. The cells were grown to high density overnight at 37°C. The next day, the preculture was used to inoculate a production culture in TB medium supplemented with 50 µg/mL kanamycin and antifoam reagent. The culture was grown in a baffled flask at 37°C to an optical density (OD600) of 0.8, and was then induced with 0.5 mM isopropyl-β-D-1-thiogalactopyranoside. Cells were grown overnight at 20°C. Cells were harvested at 4000 × g for 20 min at 12°C, and cell pellets were resuspended in twice their volume of Buffer A (50 mM 4-(2-hydroxyethyl)-1-piperazineethanesulfonic acid (HEPES) pH 7.5, 500 mM KCl) with 5 mM MgCl2 and DNase I (Roche, Basel, Switzerland). Cells were lysed by sonication using a SonoplusGM200 (BANDELIN electronic GmbH & Co. KG, Berlin, Germany) equipped with a KE76 tip at 50% amplitude for 3× 1 min of 1 s on/off pulses with 1 min pause between each cycle. The lysates were cleared by centrifugation at 100,000 × g for 1 h at 8°C, and the supernatant was then filtered through 0.45 µm filters (Sarstedt, Nümbrecht, Germany). For affinity purification, an Äkta Start FPLC system (formerly GE Healthcare, now Cytiva, Marlborough, MA, USA) with two stacked 1 mL Ni-NTA columns (HiTrap HP, Cytiva) was used. The clarified lysate was applied to the columns, which were equilibrated with Buffer A. The column was washed with Buffer A + 75 mM imidazole and eluted with Buffer A + 500 mM imidazole. The eluate was desalted using two stacked 5 mL HiTrap desalting columns (Sephadex G-25 resin, Cytiva) and protein elution buffer (25 mM Tris-HCl pH 7.4, 100 mM NaCl). Protein concentration was calculated from absorbance at 280 nm (A280) on a NanoDrop2000 with extinction coefficients calculated using ProtParam (https://web.expasy.org/protparam/). Purified TetR was aliquoted, snap-frozen in liquid nitrogen and stored at -70°C.

### *In vitro* transcription-translation (IVTT) and transcription-translation coupled DNA replication (TTcDR) assays

Our standard IVTT and TTcDR reactions, unless otherwise noted, were set up by adding the following components at their final concentrations according to the PURE manufacturer’s instructions: 1x PURE energy mix (from commercial source or modified as previously described (9, 10), 1x PURE proteins, 1x PURE ribosomes, 1 U/µL murine RNase inhibitor (New England Biolabs, catalog no.: M0314S), 4 nM of linear DNA template and different concentrations of plasmid DNA template (indicated if not equimolar). 16x linear-to-circular DNA ratio refer to 0.25 nM plasmid DNA and 4 nM linear DNA. TTcDR-compatible EMs were prepared at 7x concentration and tRNAs, amino acids, and rNTPs were added separately. IVTT reactions expressing the Pepper aptamer contained 10 µM HBC620 (MedChemExpress, Monmouth Junction, NJ, USA, catalog no.: HY-133520). TTcDR reactions contained 2.4 mM dNTPs (0.6 mM of each dNTP). The reaction compositions are shown in details in Table S5. We have also provided a pipetting scheme as an Excel sheet (Data Availability). The sample volume of an IVTT/TTcDR reaction was 10 µL and was performed in replicates of n = 3. The assumed TTcDR conditions based on PUREfrex 1.0 (48) with the Mutschler energy mix (10) or Ichihashi energy mix (9) are shown in Table S6.

For IVTT measurement, samples were mixed well by pipetting and 3x 10 µL were immediately transferred to a 384-well, small-volume, black, optically clear, flat-bottomed, medium-binding microtiter plate (Greiner Bio-One, Kremsmünster, Austria; catalog no.: 788096). Plates were centrifuged for 30 s in a small benchtop plate centrifuge (VWR, Radnor, PA, USA) prior to measurement. Reactions were characterized in triplicate on a plate reader (Infinite M200, Tecan, Männedorf, Switzerland) at 30°C with 30 s shaking before each fluorescence reading at the following excitation and emission wavelengths (Ex/Em): sfGFP: 485/528 nm; Pepper: 575/620 nm; Azurite: 383/447 nm. Readings from the bottom of the plate provide more accurate measurements than readings from the top.

For TTcDR measurement, samples were mixed well by pipetting and 3x 10 µL were transferred into 8-well PCR strips. 2 µL of each sample (0 h time point) was quenched in 18 µL quench solution (25 mM Tris-HCl pH 7.9, 0.11% Triton X-100, 1.65 mM EDTA). The remaining samples were incubated at 30°C. After 8 hours (unless stated differently) 2 µL of each sample (end time point) was quenched in 18 µL quench solution. Quenched samples were well mixed by pipetting and stored at -20°C.

### qPCR assays and analysis

The quenched TTcDR samples were mixed well by pipetting and diluted 200-fold in two steps (1) 2 µL sample + 38 µL analysis solution (25 mM Tris-HCl pH 7.9, 0.105% Triton X-100); 2) 2 µL diluted sample + 18 µL analysis solution). 2 µL of each sample were assayed in triplicate by qPCR (10 µL reactions) using Luna Universal qPCR Master Mix (New England Biolabs, catalog no.: M3003X) with oligonucleotides oMHB010+11 and the following cycling protocol: 1x 50°C for 1 min, 1x 95°C for 1 min, 30x (95°C for 0:15 min, 60°C for 30 s, fluorescence reading) + final melting curve (slow increase of 0.075°C/s from 60°C to 95°C with frequent fluorescence reading). TTcDR samples were diluted a total of 10,000-fold from the TTcDR reaction to the qPCR reaction.

We used the LinReg PCR tool (https://www.gear-genomics.com/rdml-tools/linregpcr.html) to linearly regress our qPCR curves and used the resulting efficiency-corrected target quantity values (N0) to calculate DNA amplification (f(t) = N0(8 h)/N0(0 h)) (60).

We performed TTcDR controls with an inactive Φ29 DNA polymerase mutant (D249E, (56)) for baseline subtraction of DNA amplification when background amplification was comparable between samples. Note that this was not the case when testing the three commercial PURE systems in Figure S1.

### Nuclease detection assays

We used a DNase detection kit (Jena Bioscience, catalog no.: PP-410S) and a RNase + DNase detection kit (Jena Bioscience, catalog no.: PP-409S, used for RNase detection only) to measure DNase and RNase contamination of commercial PURE systems and self-produced EMs. Therefore, we set up PURE reactions according to the PURE manufacturer’s instructions, mixed them 1:1 with the (RNase +) DNase Detection Master Mix, and measured the nuclease contamination in a qPCR cycler according to the respective nuclease detection kit instructions. Nuclease activities were standardized using the nuclease standards provided with the kits.

### Statistical Analysis

All data presented, unless otherwise noted, are the mean of n = 3 technical replicates ± (error-propagated) standard deviation (SD).

### Web Sites

The LinReg PCR server was used for linear regression of qPCR data (https://www.gear-genomics.com/rdml-tools/linregpcr.html, https://www.gear-genomics.com/rdml-tools/tableshaper.html) (60). ProtParam was used to calculate the molecular weight and theoretical extinction coefficients of all proteins used in this study (https://web.expasy.org/protparam/) (61).

## Data Availability

The data and DNA sequences underlying this article, as well as an Excel sheet for setting up TTcDR experiments, are available in FigShare at dx.doi.org/10.6084/m9.figshare.27315201. The following plasmids have been deposited at Addgene (pTE5415, pTE5422, pTE5425-5430, pTE5444, pTE5448, pTE5451, pTE5452) and are accessible under catalogue numbers #229030-229037, 231064-231067.

## Acknowledgments

The authors thank Nitin Bohra, Scott Scholz, Franziska Otto, René Inckemann, Markus Meier, and Blake Rasor for helpful discussions. We also thank Zhanar Abil and Christophe Danelon for helpful discussions and for providing plasmid G95, encoding the *p2* wild-type gene. Special thanks go to Nitin Bohra for general help with the PURE system and purification of tRNAs, and to Luca Brenker for help with the Pepper aptamer.

## Author Contributions

Conceptualization, S.B., and T.J.E.; Methodology, S.B., M.H.B. and I.G.K.; Investigation, S.B. and M.H.B.; Visualization: S.B. and M.H.B.; Writing – Original Draft, S.B.; Writing – Review & Editing, S.B. and T.J.E.; Funding Acquisition, T.J.E.; Resources, T.J.E.; Supervision, S.B. and T.J.E.

## Author ORCIDs

