## Supplementary Information for "Genetically encoded control of *in vitro* transcription-translation coupled DNA replication"

**Supplementary Note S3:** Effect of chaperone mixes and EF-P on circuit performance. ... 5

#### Supplementary Notes

**Supplementary Note S1:** Effect of DNA topology on gene expression under standard TTcDR conditions.

We observed reduced performance of the TetR circuit with IVTT output under our standard TTcDR conditions, compared to PUREfrex 1.0 conditions (Figure 1F, S5F). We further observed that at equimolar concentration of linear DNA, encoding Φ29 DNA polymerase, and circular DNA, encoding EilR (no repression condition), plasmid DNA was replicated 4-fold, while the self-replication scheme showed >20-fold (Figure 2B, S11). We hypothesized that a template expression bias might be at work under TTcDR conditions, which reduces the production of reporter protein encoded on linear DNA templates.

To elucidate these observations, we first compared the effect of DNA topology on transcriptional (Pepper aptamer) and translational (sfGFP) level in PUREfrex 1.0 and under TTcDR conditions (Figure S6A). On transcriptional level, we observed faster rates and higher outputs from plasmid DNA in both conditions, but more pronounced under standard TTcDR conditions. On translational level, the DNA topology had no effect on the rate in PUREfrex 1.0, but under standard TTcDR conditions, the use of circular DNA approximately doubled rate and output. Taken together these results indicate a preference for plasmid DNA templates under standard TTcDR conditions.

We next sought to elucidate potential competing effects between linear and circular DNA templates. We used a fluorescent three-color readout to measure transcriptional and translational output (sfGFP-Pepper) from one template and translational output (Azurite, blue channel) from a competing template (Figure 6B,D). We tested the effect of DNA topology (linear template versus plasmid template) in the presence of competing DNA at an equimolar concentration and normalized the data to the corresponding output without competing DNA. While in PUREfrex 1.0 the topology of the competing DNA had no effect, the relative output, transcriptional and translational, was always higher when competing with a linear DNA, indicating stronger competition from circular DNA templates. For example, under TTcDR conditions, *in vitro* transcription of the Pepper aptamer from linear DNA was 3-fold higher in the presence of linear compared to circular competing DNA. Strikingly, at equimolar concentrations, ~30% Pepper aptamer and ~20% sfGFP were produced from linear DNA, but >90% Azurite from circular DNA (all numbers relative to the respective template alone). To shift expression from plasmid-encoded reporter to the reporter on the linear template, we next titrated the ratio of linear to circular DNA. As the ratio of linear DNA increased, we observed a higher output of sfGFP and Pepper, while Azurite production decreased (Figure S6C).

These results show that under standard TTcDR conditions, plasmid DNA is preferred over linear DNA templates. These findings need to be considered in the construction of genetic circuits. We showed that titrating the linear-to-circular DNA ratio is an applicable measure to balance the protein output from circular and linear DNA templates to improve IVTT and TTcDR output and genetic circuit performance (Figure S6F, Figure 2B).

**Supplementary Note S2:** Effect of encoding the genetic circuit on a single plasmid.

Our circuit experiments with IVTT output allowed us to monitor transcriptional and translational reporter formation with great time resolution. We observed that in the beginning of the reaction, when only low concentrations of TetR have been produced, the system is not well repressed, resulting in about 4-fold reduced circuit repression when co-producing TetR compared to supplementing 500 nM TetR (Figure 1F). This points out that improving circuit repression in the early phase of the circuit promises to drastically improve the overall circuit performance. Our genetic circuit is based on two reporter template molecules (one linear and one circular DNA molecule) from which proteins are produced with interact with the other DNA molecule, respectively, so that protein-DNA interactions are intermolecular. The PURE system is highly diluted compared to the cellular environment of *Escherichia coli*. Consequently, in an intermolecular circuit, the produced aTF and DNA polymerase must diffuse longer distances to another DNA molecule to exert their function (Figure S8A). To further improve the genetic circuit, we hypothesized that encoding all genes of the circuit on the same plasmid and co-localizing the transcription-translation machinery at the corresponding DNA molecule could increase the performance of the circuit (1). We reasoned that this 'intramolecular' setup would also circumvent the observed competition between circular and linear DNA templates (Figure S6, Note S1).

However, we observed increased leakage of the intramolecular TetR circuit and reduced DNA replication of the non-repressing condition (EilR; Figure S8B-D). Downregulation of Φ29 DNA polymerase production using a ribosome binding site (RBS) with lower translation initiation strength to increase the ratio of TetR to DNA polymerase further reduced DNA replication without increasing repression. Overall, we observed the best performance of the intramolecular circuit with the previously used RBS B0035 at 3.3-fold repression and 1.5-fold induction, indicating that the intramolecular circuit was not able to compete with the intermolecular approach.

**Supplementary Note S3:** Effect of chaperone mixes and EF-P on circuit performance.

Protein folding in bacteria is assisted by chaperons, such as the DnaKJE machinery and GroEL/ES (2), which also significantly improved folding and solubility of proteins produced in the PURE system (3). Furthermore, protein synthesis in the PURE system yields truncated products (4–6). Especially consecutive proline stretches may lead to stalling but can be rescued by elongation factor P (EF-P) without peptide truncation (7, 8). EF-P has been shown to improve full-length protein translation in the PURE system (4, 8) and has recently been essential to regenerate 20 aminoacyl-tRNA synthetases in the PURE system (9). GeneFrontier, the commercial provider of PUREfrex 1.0, offers chaperone mixes of the DnaKJE machinery (#PF003) and GroEL/ES (#PF004), as well as EF-P (#PFS052).

We hypothesized that chaperone mixes as well as EF-P may improve TTcDR and genetic circuit performance. Improved Φ29 DNA polymerase wild-type (69 kDa enzyme) synthesis may result in improved DNA replication, while improved TetR synthesis (contains a PP site in its effector binding domain) may result in improved repression and induction, if TetR is truncated or not correctly folded in the PURE system, which may result in the loss of allostery.

We observed 3.6-fold improved DNA replication when supplementing GroE mix, while the DnaK mix reduced DNA replication by 5.2-fold. EF-P supplementation had no significant effect. Note that GroE supplementation also slightly improved repression and induction of the TetR circuit (Figure S10). Nevertheless, we have not observed significant improvements resolving the poor induction of the system, e.g. by improving TetR folding and, if TetR is misfolded, restoring its allostery.

### Supplementary Tables

**Supplementary Table S1:** All strains used in this study.

| Strain ID | Background strain | Plasmid & resistance marker# | Use | Reference |
|---|---|---|---|---|
| sTetR | *Escherichia coli* BL21 (DE3) | pJBL701, $Kan^R$ | Protein production of TetR | Jung et al., (2020) (10) |

### $Kan^R$, kanamycin resistance

| Commercial strain | Genotype | Source |
|---|---|---|
| *Escherichia coli* NEB 5-alpha | *fhuA2Δ(argF-lacZ)U169 phoA glnV44 Φ80Δ(lacZ)M15 gyrA96 recA1 relA1 endA1 thi-1 hsdR17* | New England Biolabs (Frankfurt am Main, Germany) |
| *Escherichia coli* BL21 (DE3) | *fhuA2 [lon] ompT gal (λ DE3) [dcm] ∆hsdS λ DE3 = λ sBamHIo ∆EcoRI-B int::(lacI::PlacUV5::T7 gene1) i21 ∆nin5* | New England Biolabs (Frankfurt am Main, Germany) |

**Supplementary Table S2:** All plasmids used in this study.

| Plasmid | Plasmid ID | Relevant features# | Reference | Used in |
|---|---|---|---|---|
| pDNAP_wt | pTE5415 | Expression vector for 10xHis-phi29 DNA polymerase (wild-type): *$P_{T7}$-lacO-g10leader-10xHis-p2 wt-T7 terminator*, $Cam^R$ | This study. Addgene: #229030 | Fig. 1D Fig. S1, S2, S4C |
| pGFP-Pepper | pTE5422 | sfGFP-Pepper reporter: *$P_{T7}$-tetO-B0035-sfGFP-pepper-T7 terminator*, $Amp^R$ | This study. Addgene: #229031 | Fig. S6, S9 |
| pDNAP_dead | pTE5425 | Expression vector for 10xHis-phi29 DNA polymerase (inactive mutant D249E): *$P_{T7}$-lacO-g10leader-10xHis-p2 dead-T7 terminator*, $Cam^R$ | This study. Addgene: #229032 | Fig. 1D Fig. S1, S2, S4C |
| pDNAP_wt-tetO | pTE5426 | *tetO*-phi29 DNA polymerase wild-type reporter: *$P_{T7}$-tetO-B0035-p2 wt-T7 terminator*, $Amp^R$ | This study. Addgene: #229033 | linear template preparation only |
| pDNAP_dead-tetO | pTE5427 | Inactive *tetO*-phi29 DNA polymerase 'dead' reporter: *$P_{T7}$-tetO-B0035-p2 D249E-T7 terminator*, $Amp^R$ | This study. Addgene: #229034 | linear template preparation only |
| pTetR | pTE5428 | Expression vector for TetR: *$P_{T7}$-lacO-B0035-TetR-T7 terminator*, $Amp^R$ | This study. Addgene: #229035 | Fig. 1F, 2B,C,F Fig. S5, S7, S10, S11, S12 |
| pEilR | pTE5429 | Expression vector for EilR: *$P_{T7}$-lacO-B0035-EilR-T7 terminator*, $Amp^R$ | This study. Addgene: #229036 | Fig. 1F, 2 Fig. S5, S7, S10, S11, S12 |
| pAzurite | pTE5430 | Azurite reporter: *$P_{T7}$-tetO-B0035-Azurite-T7 terminator*, $Amp^R$ | This study. Addgene: #229037 | Fig. S6 |
| pTetR-tetO-B0035-DNAP_wt | pTE5431 | Vector encoding *$P_{T7}$-lacO-B0035-**TetR**-T7 terminator* (reverse direction) and *$P_{T7}$-tetO-**B0035**-p2 **wt**-T7 terminator*, $Amp^R$ | This study. | Fig. S8 |
| pTetR-tetO-B0035-DNAP_dead | pTE5432 | Vector encoding *$P_{T7}$-lacO-B0035-**TetR**-T7 terminator* (reverse direction) and *$P_{T7}$-tetO-**B0035**-p2 **dead**-T7 terminator*, $Amp^R$ | This study. | Fig. S8 |
| pEilR-tetO-B0035-DNAP_wt | pTE5433 | Vector encoding *$P_{T7}$-lacO-B0035-**EilR**-T7 terminator* (reverse direction) and *$P_{T7}$-tetO-**B0035**-p2 **wt**-T7 terminator*, $Amp^R$ | This study. | Fig. S8 |

| | | | | |
|---|---|---|---|---|
| pEilR-tetO-B0035-DNAP_dead | pTE5434 | Vector encoding $P_{T7}$-*lacO-B0035*-***EilR***-*T7 terminator* (reverse direction) and $P_{T7}$-*tetO*-***B0035***-*p2* ***dead***-*T7 terminator*, Amp$^R$ | This study. | Fig. S8 |
| pTetR-tetO-B0032-DNAP_wt | pTE5435 | Vector encoding $P_{T7}$-*lacO-B0035*-***TetR***-*T7 terminator* (reverse direction) and $P_{T7}$-*tetO*-***B0032***-*p2* ***wt***-*T7 terminator*, Amp$^R$ | This study. | Fig. S8 |
| pTetR-tetO-B0034-DNAP_wt | pTE5436 | Vector encoding $P_{T7}$-*lacO-B0035*-***TetR***-*T7 terminator* (reverse direction) and $P_{T7}$-*tetO*-***B0034***-*p2* ***wt***-*T7 terminator*, Amp$^R$ | This study. | Fig. S8 |
| pTetR-tetO-B0064-DNAP_wt | pTE5437 | Vector encoding $P_{T7}$-*lacO-B0035*-***TetR***-*T7 terminator* (reverse direction) and $P_{T7}$-*tetO*-***B0064***-*p2* ***wt***-*T7 terminator*, Amp$^R$ | This study. | Fig. S8 |
| pEilR-tetO-B0032-DNAP_wt | pTE5438 | Vector encoding $P_{T7}$-*lacO-B0035*-***EilR***-*T7 terminator* (reverse direction) and $P_{T7}$-*tetO*-***B0032***-*p2* ***wt***-*T7 terminator*, Amp$^R$ | This study. | Fig. S8 |
| pEilR-tetO-B0034-DNAP_wt | pTE5439 | Vector encoding $P_{T7}$-*lacO-B0035*-***EilR***-*T7 terminator* (reverse direction) and $P_{T7}$-*tetO*-***B0034***-*p2* ***wt***-*T7 terminator*, Amp$^R$ | This study. | Fig. S8 |
| pEilR-tetO-B0064-DNAP_wt | pTE5440 | Vector encoding $P_{T7}$-*lacO-B0035*-***EilR***-*T7 terminator* (reverse direction) and $P_{T7}$-*tetO*-***B0064***-*p2* ***wt***-*T7 terminator*, Amp$^R$ | This study. | Fig. S8 |
| pDNAP_wt-noHisTag | pTE5441 | Expression vector for phi29 DNA polymerase (wild-type): $P_{T7}$-*lacO-g10leader-p2 wt-T7 terminator*, Cam$^R$ | This study. | linear template preparation only |
| pDNAP_ Sakatani | pTE5442 | Expression vector for 10x-phi29 DNA polymerase (Sakatani): $P_{T7}$-*lacO-g10leader-10xHis-p2 sak-T7 terminator*, Cam$^R$ | This study. | linear template preparation only |
| pDNAP_ Povilaitis | pTE5443 | Expression vector for 10x-phi29 DNA polymerase (Poviliatis): $P_{T7}$-*lacO-g10leader-10xHis-p2 pov-T7 terminator*, Cam$^R$ | This study. | linear template preparation only |
| pDNAP_ PovSak | pTE5444 | Expression vector for 10x-phi29 DNA polymerase (PovSak): $P_{T7}$-*lacO-g10leader-10xHis-p2 povsak-T7 terminator*, Cam$^R$ | This study. Addgene: #231064 | linear template preparation only |
| pDNAP_Gao | pTE5445 | Expression vector for 10x-phi29 DNA polymerase (Gao): $P_{T7}$-*lacO-g10leader-10xHis-p2 gao-T7 terminator*, Cam$^R$ | This study. | linear template preparation only |
| pDNAP_ Okauchi | pTE5446 | Expression vector for 10x-phi29 DNA polymerase (Okauchi): $P_{T7}$-*lacO-g10leader-10xHis-p2 ok-T7 terminator*, Cam$^R$ | This study. | linear template preparation only |
| pDNAP_ SakOk | pTE5447 | Expression vector for 10x-phi29 DNA polymerase (SakOk): $P_{T7}$-*lacO-g10leader-10xHis-p2 sakok-T7 terminator*, Cam$^R$ | This study. | linear template preparation only |
| pDNAP_ PovSakOk | pTE5448 | Expression vector for 10x-phi29 DNA polymerase (PovSakOk): $P_{T7}$-*lacO-g10leader-10xHis-p2 povsakok-T7 terminator*, Cam$^R$ | This study. Addgene: #231065 | linear template preparation only |
| pDNAP_PovSak-tetO | pTE5449 | *tetO*-phi29 DNA polymerase PovSak reporter: $P_{T7}$-*tetO-B0035-p2 povsak-T7 terminator*, Amp$^R$ | This study. | linear template preparation only |
| pDNAP_wt-tetO-HisTag | pTE5450 | *tetO*-10xHis-phi29 DNA polymerase wild-type reporter: $P_{T7}$-*tetO-B0035-10xHis-p2 wt-T7 terminator*, Amp$^R$ | This study. | linear template preparation only |
| pDNAP_PovSak-tetO-HisTag | pTE5451 | *tetO*-10xHis-phi29 DNA polymerase PovSak reporter: $P_{T7}$-*tetO-B0035-10xHis-p2 povsak-T7 terminator*, Amp$^R$ | This study. Addgene: #231066 | linear template preparation only |
| pDNAP_dead-tetO-HisTag | pTE5452 | Inactive *tetO*-10xHis-phi29 DNA polymerase wild-type reporter: $P_{T7}$-*tetO-B0035-10xHis-p2 dead-T7 terminator*, Amp$^R$ | This study. Addgene: #231067 | linear template preparation only |

| | | | |
|---|---|---|---|
| pJBL701 | Expression vector for C-terminally His-tagged TetR from *Escherichia coli*, $Kan^R$ | Jung et al., 2020 (10) Addgene: #140371 | TetR production & purification |

[#] $Amp^R$, ampicillin resistance; $Cam^R$, chloramphenicol resistance; $Kan^R$, kanamycin resistance
All sequences are provided as GenBank files on FigShare at dx.doi.org/10.6084/m9.figshare.27315201.

**Supplementary Table S3:** All linear DNA templates used in this study.

| Template ID | Description | PCR template | Oligonucleotides | Used in |
|---|---|---|---|---|
| tGFP-Pepper | sfGFP-Pepper reporter: *$P_{T7}$-tetO-B0035-sfGFP-pepper-T7 terminator* | pTE5422 | oSB0021+22 | Fig. 1D,F<br>Fig. S1,3-6 |
| tAzurite | Azurite reporter: *$P_{T7}$-tetO-B0035-Azurite-T7 terminator* | pTE5430 | oSB0021+22 | Fig. S6 |
| tDNAP_wt | phi29 DNA polymerase wild-type reporter: *$P_{T7}$-lacO-g10leader-10xHis-p2 wt-T7 terminator* | pTE5415 | oSB0118+20 | Fig. 2E<br>Fig. S9 |
| tDNAP_dead | phi29 DNA polymerase dead reporter: *$P_{T7}$-lacO-g10leader-10xHis-p2 dead-T7 terminator* | pTE5425 | oSB0118+20 | Fig. 2E<br>Fig. S9 |
| tDNAP_wt-tetO | *tetO*-phi29 DNA polymerase wild-type reporter: *$P_{T7}$-tetO-B0035-p2 wt-T7 terminator* | pTE5426 | oSB0021+22 | Fig. 2B,C,F<br>Fig. S7, S9-12 |
| tDNAP_dead-tetO | Inactive *tetO*-phi29 DNA polymerase 'dead' reporter: *$P_{T7}$-tetO-B0035-p2 D249E-T7 terminator* | pTE5427 | oSB0021+22 | Fig. 2B,C,F<br>Fig. S7, S9-12 |
| tDNAP_wt-noHisTag | phi29 DNA polymerase wt reporter: *$P_{T7}$-lacO-g10leader-p2 wt-T7 terminator* | pTE5441 | oSB0118+20 | Fig. 2E |
| tDNAP_ Sakatani | phi29 DNA polymerase Sakatani reporter: *$P_{T7}$-lacO-g10leader-10xHis-p2 sak-T7 terminator* | pTE5442 | oSB0118+20 | Fig. 2E |
| tDNAP_ Povilaitis | phi29 DNA polymerase Povilaitis reporter: *$P_{T7}$-lacO-g10leader-10xHis-p2 pov-T7 terminator* | pTE5443 | oSB0118+20 | Fig. 2E |
| tDNAP_ PovSak | phi29 DNA polymerase PovSak reporter: *$P_{T7}$-lacO-g10leader-10xHis-p2 povsak-T7 terminator* | pTE5444 | oSB0118+20 | Fig. 2E<br>Fig. S9 |
| tDNAP_Gao | phi29 DNA polymerase Gao reporter: *$P_{T7}$-lacO-g10leader-10xHis-p2 gao-T7 terminator* | pTE5445 | oSB0118+20 | Fig. 2E |
| tDNAP_ Okauchi | phi29 DNA polymerase Okauchi reporter: *$P_{T7}$-lacO-g10leader-10xHis-p2 ok-T7 terminator* | pTE5446 | oSB0118+20 | Fig. 2E |
| tDNAP_ SakOk | phi29 DNA polymerase SakOk reporter: *$P_{T7}$-lacO-g10leader-10xHis-p2 sakok-T7 terminator* | pTE5447 | oSB0118+20 | Fig. 2E |
| tDNAP_ PovSakOk | phi29 DNA polymerase PovSakOk reporter: *$P_{T7}$-lacO-g10leader-10xHis-p2 povsakok-T7 terminator* | pTE5448 | oSB0118+20 | Fig. 2E |
| tDNAP_PovSak-tetO | *tetO*-phi29 DNA polymerase PovSak reporter: *$P_{T7}$-tetO-B0035-p2 povsak-T7 terminator* | pTE5449 | oSB0021+22 | Fig. S9 |

| | | | | |
|---|---|---|---|---|
| tDNAP_wt-tetO-HisTag | *tetO*-10xHis-phi29 DNA polymerase wild-type reporter: *$P_{T7}$-tetO-B0035-10xHis-p2 wt-T7 terminator* | pTE5450 | oSB0021+22 | Fig. S9 |
| tDNAP_PovSak-tetO-HisTag | *tetO*-10xHis-phi29 DNA polymerase PovSak reporter: *$P_{T7}$-tetO-B0035-10xHis-p2 povsak-T7 terminator* | pTE5451 | oSB0021+22 | Fig. S2F<br>Fig. S9 |
| tDNAP_dead-tetO-HisTag | Inactive *tetO*-10xHis-phi29 DNA polymerase wild-type reporter: *$P_{T7}$-tetO-B0035-10xHis-p2 dead-T7 terminator* | pTE5452 | oSB0021+22 | Fig. S9 |

**Supplementary Table S4:** All oligonucleotides used for preparation of linear DNA templates and qPCR.

| **Oligo ID** | **Sequence [5' to 3']** | **Used for** |
|---|---|---|
| oMHB0010 | GGTGTAGGTCGTTCGCTCCAA | qPCR |
| oMHB0011 | CCTCGCTCTGCTAATCCTGTTACC | qPCR |
| oSB0020 | GAAGCCTGCATAACGCGAAG | PCR amplification of linear DNA templates |
| oSB0021 | CGGTTCCTGGCCTTTTGC | PCR amplification of linear DNA templates |
| oSB0022 | GATAGGTGCCTCACTGATTAAGC | PCR amplification of linear DNA templates |
| oSB0118 | ATGCGTCCGGCGTAG | PCR amplification of linear DNA templates |

**Supplementary Table S5:** IVTT and TTcDR reaction composition based on commercial PURE systems.

| Variant | Reagent | Stock concentration | Final concentration | Remarks |
|---|---|---|---|---|
| Mutschler EM | Customized EM (11) | 700 mM HEPES KOH, pH 8.0<br>490 mM potassium glutamate<br>2.625 mM spermidine<br>175 mM creatine phosphate<br>55.3 mM hemi-magnesium glutamate<br>42 mM dithiothreitol (DTT) | 100 mM HEPES KOH, pH 8.0<br>70 mM potassium glutamate<br>0.375 mM spermidine<br>25 mM creatine phosphate<br>7.9 mM hemi-magnesium glutamate<br>6 mM DTT | 7x stock |
| | Commercial EM | PUREfrex 1.0 solution I: 2x<br>PUREfrex 2.0 solution I: 2x<br>PURExpress solution A: 2.5x | 0.1x | PUREfrex 1.0 EM (solution I) composition: (12) |
| Ichihashi EM | Customized EM (13) | 700 mM HEPES KOH, pH 7.6<br>490 mM potassium glutamate<br>2.625 mM spermidine<br>175 mM creatine phosphate<br>55.3 mM magnesium acetate<br>42 mM DTT<br>70 ng/µL 10-formyl-5,6,7,8-tetrahydrofolic acid (THF) | 100 mM HEPES KOH, pH 7.6<br>70 mM potassium glutamate<br>0.375 mM spermidine<br>25 mM creatine phosphate<br>7.9 mM magnesium acetate<br>6 mM DTT<br>10 ng/µL THF | 7x stock |
| General | Amino acids | 6 mM | 0.36 mM | 16.67x stock |
| | dNTPs | 100 mM total<br>(25 mM each) | 2.4 mM total<br>(0.6 mM each) | 41.67x stock |
| | rNTPs | 18.75 mM ATP, 12.5 mM GTP<br>6.25 mM UTP/CTP | 0.375 mM ATP, 0.25 mM GTP<br>0.125 mM UTP/CTP | 50x stock |
| | tRNA stock | 53.3 g/L | 0.52 g/L | 102.5x stock |
| | Murine RNase inhibitor | 40 U/µL | 1 U/µL | 40x stock |
| | PURE proteins & ribosomes | PUREfrex 1.0: 20x solution II (proteins), 20x solution III (ribosomes)<br>PUREfrex 2.0: 20x solution II (proteins), 10x solution III (ribosomes)<br>PURExpress: 3.33x solution B (combined) | 1x | PUREfrex 1.0 EM composition: (12) |
| | HBC620 | 1 mM | 10 µM | **in IVTT reactions only** |
| | DNA | 40 nM | 4 nM | unless otherwise noted |
| | Anhydro-tetracycline | 250 µM | 12.5 µM | 50 µM shows inhibition |
| | TetR (purified) | 5 µM | 500 nM | in supplemented conditions only |
| | aTFs, ligands, etc. | various | various | individual |

See manufacturer's instruction for the composition of the commercial PURE systems tested: Gene Frontier: PUREfrex 1.0 & 2.0; New England Biolabs: PURExpress.

**Supplementary Table S6:** Assumed TTcDR conditions based on PUREfrex 1.0 and the Mutschler EM or the Ichihashi EM.

| Reagent | **PUREfrex 1.0** (12) | **Mutschler EM** (11) | | **Ichihashi EM** (13) |
|---|---|---|---|---|
| | **1x EM, commercial** | **1x EM, customized** | **1x EM + 0.1x PUREfrex 1.0 EM** | **1x EM customized** |
| HEPES KOH, pH 7.6 | 50 mM | - | 5 mM | 100 mM |
| HEPES KOH, pH 8.0 | - | 100 mM | 100 mM | - |
| potassium glutamate | 100 mM | 70 mM | 80 mM | 70 mM |
| hemi-magnesium glutamate | - | 7.9 mM | 7.9 mM | - |
| magnesium acetate | 13 mM | - | 1.3 mM | 10.5 mM |
| spermidine | 2 mM | 0.375 mM | 0.575 mM | 0.375 mM |
| DTT | 1 mM | 6 mM | 6.1 mM | 6 mM |
| 20 amino acids, ea. | 0.30 mM | 0.36 mM | 0.39 mM | 0.36 mM |
| THF | 0.01 g/L | - | 0.001 g/L | 0.01 g/L |
| *E. coli* tRNAs | 2.24 g/L | 0.52 g/L | 0.742 g/L | 0.52 g/L |
| ATP | 2 mM | 0.375 mM | 0.575 mM | 0.375 mM |
| GTP | 2 mM | 0.25 mM | 0.45 mM | 0.25 mM |
| CTP | 1 mM | 0.125 mM | 0.225 mM | 0.125 mM |
| UTP | 1 mM | 0.125 mM | 0.225 mM | 0.125 mM |
| creatine phosphate | 20 mM | 25 mM | 27 mM | 25 mM |
| dNTPs | - | 2.4 mM | 2.4 mM | 2.4 mM |

See Shimizu et al. for information on proteins and ribosomes (12).

**Supplementary Table S7:** Information on screened Φ29 DNA polymerase variants.

| Variant | Mutation(s)* | Selected in | Reference |
|---|---|---|---|
| Sakatani | K555T, D570N | self-replication in TTcDR | Sakatani et al. (14) |
| Povilaitis | M8R, V51A, M97T, G197D, E221K | *in vivo* self-replication | Povilaitis et al. (15) |
| Okauchi | D398N, D423N | self-replication in TTcDR | Okauchi et al. (16), evo3 |
| Gao | E375S | rational engineering | Gao et al. (17) |
| SakOk | D398N, D423N, K555T, D570N | This work. | Combination of (14, 16) |
| PovSak | M8R, V51A, M97T, G197D, E221K, K555T, D570N | This work. | Combination of (14, 15) |
| PovSakOk | M8R, V51A, M97T, G197D, E221K, D398N, D423N, K555T, D570N | This work. | Combination of (14–16) |

* counting based on the MKHM starting sequence of Φ29 DNA polymerase.

#### Supplementary Figures

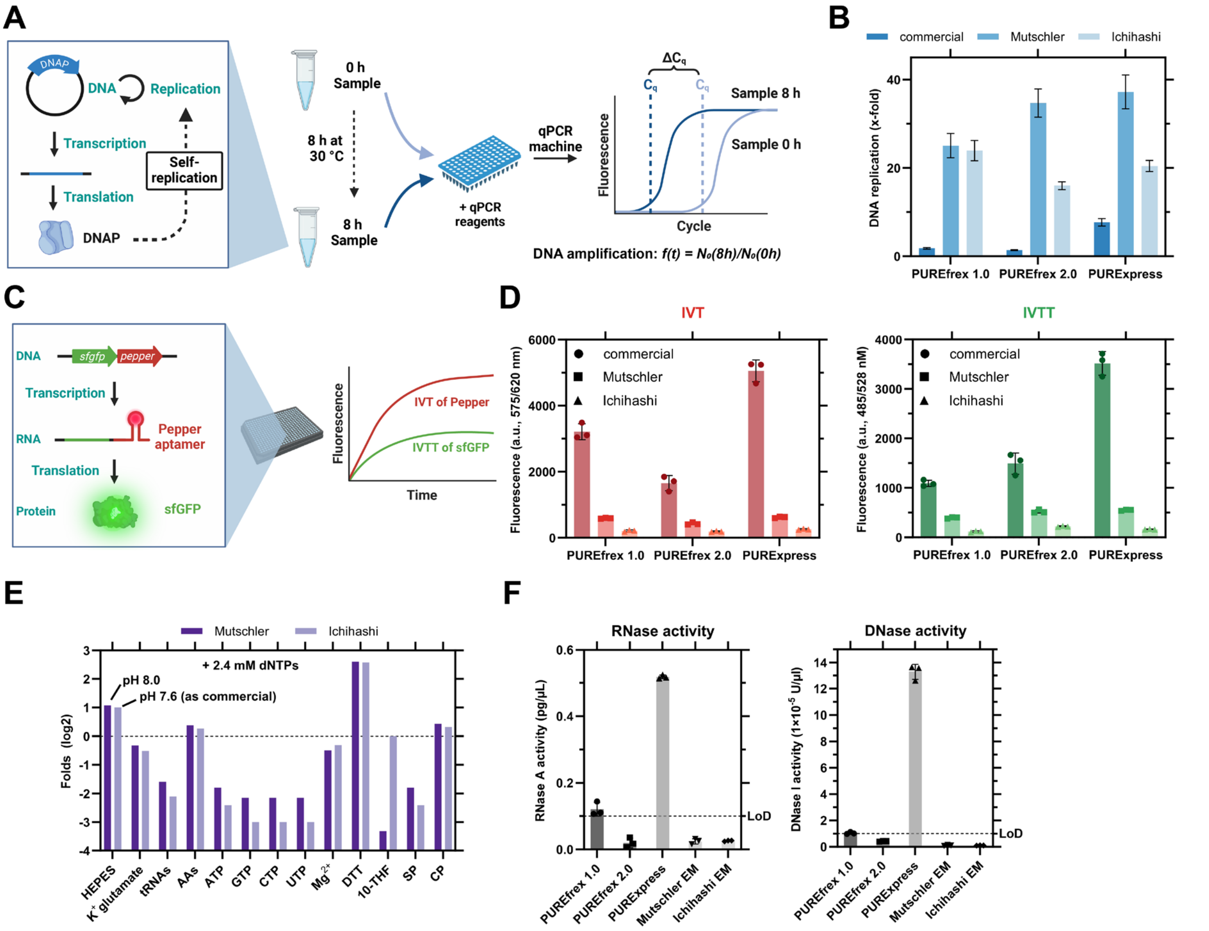

**Figure S1:** Characterization of stand-alone Φ29 DNA polymerase-based transcription-translation coupled DNA replication (TTcDR) systems prepared from three commercial PURE systems for DNA replication activity (**A,B**), transcription-translation (IVTT) capacity (**C,D**), and nuclease contamination (**F**). **A:** Schematic of TTcDR based on Φ29 DNA polymerase-driven self-replication and qPCR analysis. Φ29 DNA polymerase is produced in the IVTT system from plasmid, which in turn is self-replicated by Φ29 DNA polymerase. DNA replication is analyzed by comparing DNA quantities measured by qPCR of samples taken before and after incubation at 30°C. **B:** DNA replication of three commercial PURE systems (PUREfrex 1.0, PUREfrex 2.0 and PURExpress) using either the respective commercial energy mix (EM) or a reported TTcDR-compatible EM (11, 13) (see **E** for EM composition). Data are not baseline corrected due to differences in nuclease contamination (see **E**) and highly variable background. qPCR data are shown in Figure S2. **C:** Schematic of IVTT based on a two-color fluorescence plate reader assay using the Pepper aptamer with HBC620 ligand to measure *in vitro* transcription (IVT) in the red channel, and sfGFP to measure *in vitro* transcription-translation (IVTT) in the green channel. **D:** IVT (left) and IVTT (right) output of the three PURE systems tested with the respective commercial EM or two TTcDR-compatible EMs. IVT and IVTT time course data are shown in Figure S3. **E:** Composition of Mutschler EM (with 0.1x PUREfrex 1.0 EM) and Ichihashi EM compared to PUREfrex 1.0 EM (12). AAs: amino acids, SP: spermidine, CP: creatine phosphate. **F:** Quantification of RNase and DNase activity in the three commercial PURE systems with their respective EMs and the two TTcDR-compatible EMs. LoD: limit of detection as reported by Jena Bioscience (Catalog no.: PP-410S and PP-409S). Data are the mean of n = 3 replicates ± SD unless otherwise noted.

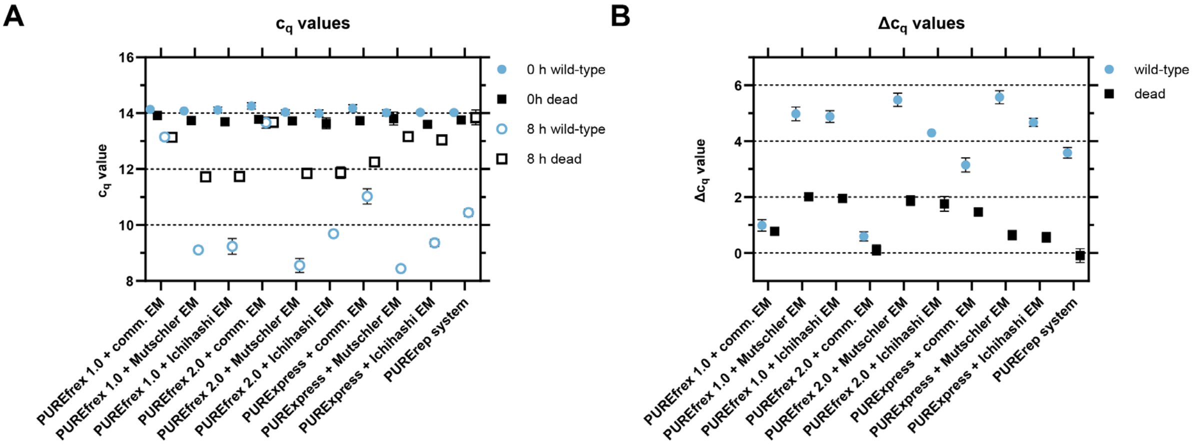

**Figure S2:** qPCR results of testing three commercial PURE systems (PUREfrex 1.0, PUREfrex 2.0 and PURExpress) with two previously reported TTcDR-compatible energy mixtures (EM) (11, 13). **A:** cq values of the experiment at time points 0 h and 8 h for wild-type and controls with an inactive Φ29 DNA polymerase 'dead' mutant D249E (18). Due to the highly variable background amplification of the inactive mutant between the different PURE system setups, we decided to show the DNA replication without background correction in Figure 1D and Figure S1B. Compare with Figure S4C for an example of background correction. **B:** Difference in $c_q$ values ($\Delta c_q$) of 0 h and 8 h time points. Data are the mean of n = 3 replicates ± SD.

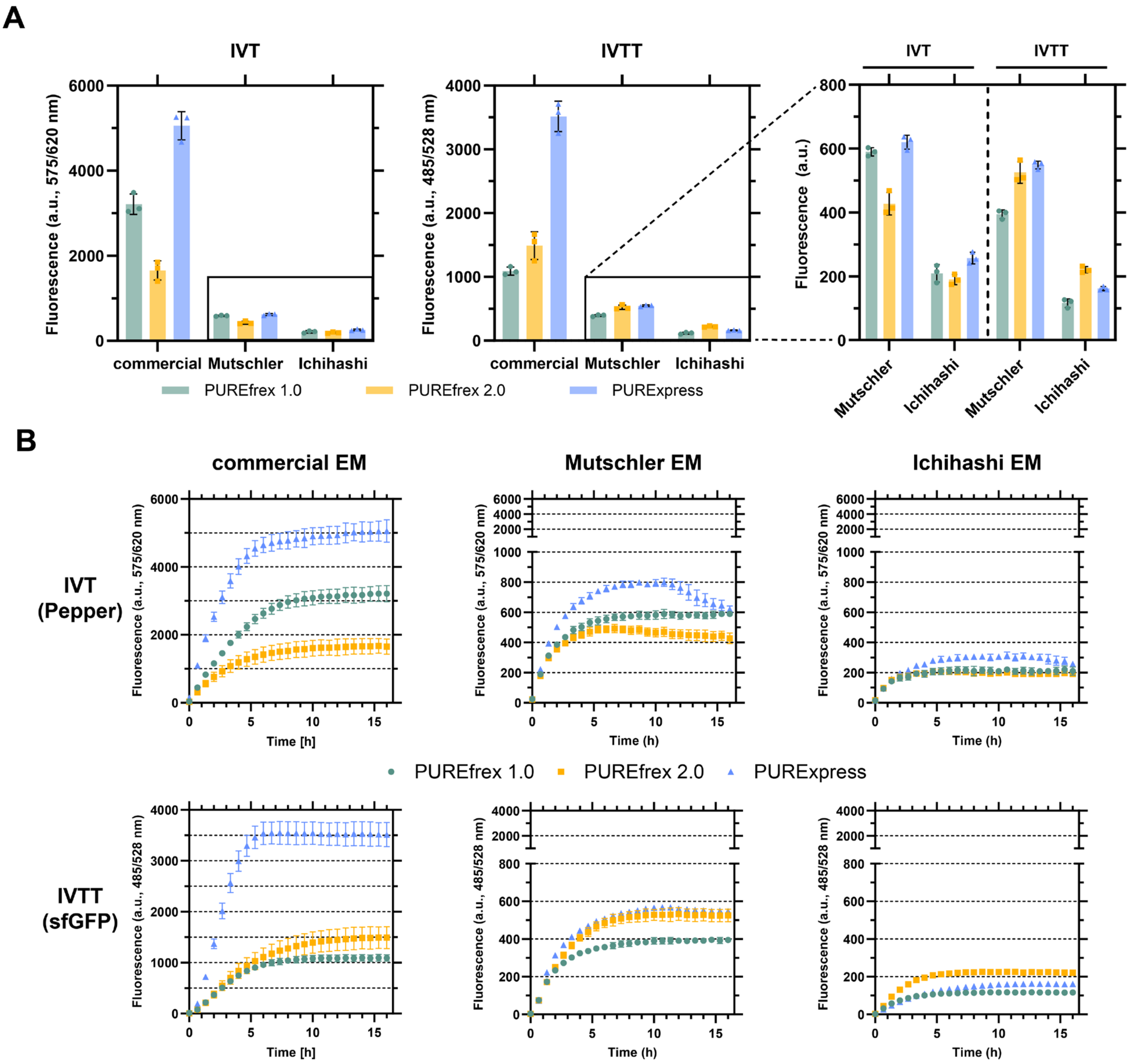

**Figure S3:** Characterization of *in vitro* transcription (IVT) and *in vitro* transcription-translation (IVTT) output of three commercial PURE systems with either their commercial EM or two previously TTcDR-compatible EMs (11, 13). **A:** 16 h time point data for IVT output (Pepper aptamer expression + HBC620 ligand, left), IVTT output (sfGFP production, center) and a close-up of IVT and IVTT output from the Mutschler and Ichihashi EMs (right). **B:** Time course data for each combination of the three PURE systems and the three EMs tested on IVT (upper panels) and IVTT (lower panels). Data are based on expression from linear DNA and are the mean of n = 3 replicates ± SD.

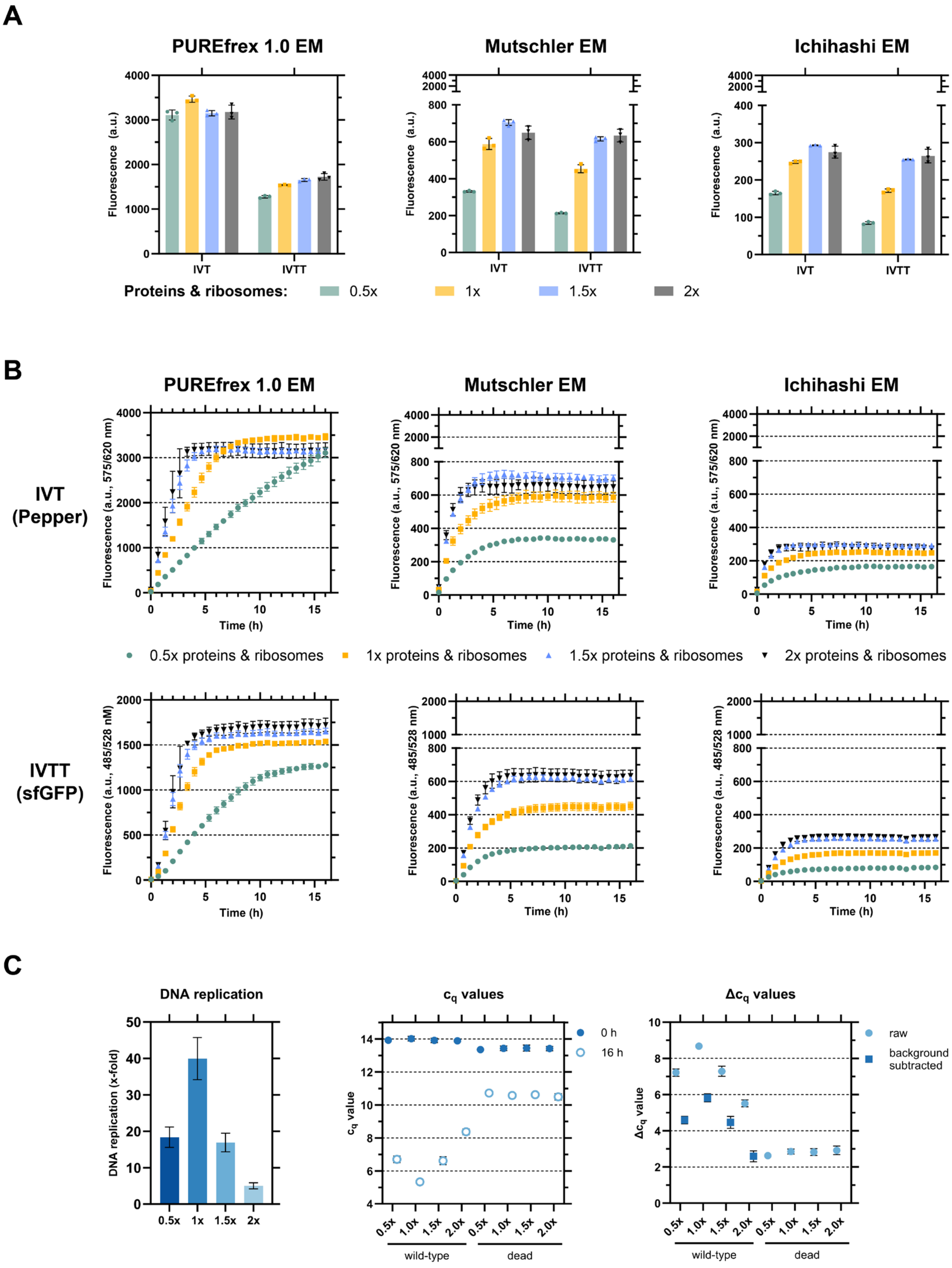

**Figure S4:** Effect of PURE protein and ribosome titration on the three commercial PURE systems in the three EMs tested. **A:** 16 h time point data for IVT output (Pepper aptamer expression + HBC620 ligand) and IVTT output (sfGFP production). **B:** Time course data for each combination of the three PURE systems and the three EMs tested on IVT (upper panels) and IVTT (lower panels). **C:** Calculated DNA replication (left) and qPCR results in $c_q$ (middle) and $\Delta c_q$ values (right) of 0 h and 16 h time points. Consistent background amplification of the inactive mutant (18) (see middle panel) allowed background subtraction of $\Delta c_q$ values. IVT and IVTT data are based on expression from linear DNA. Data are the mean of n = 3 replicates ± SD.

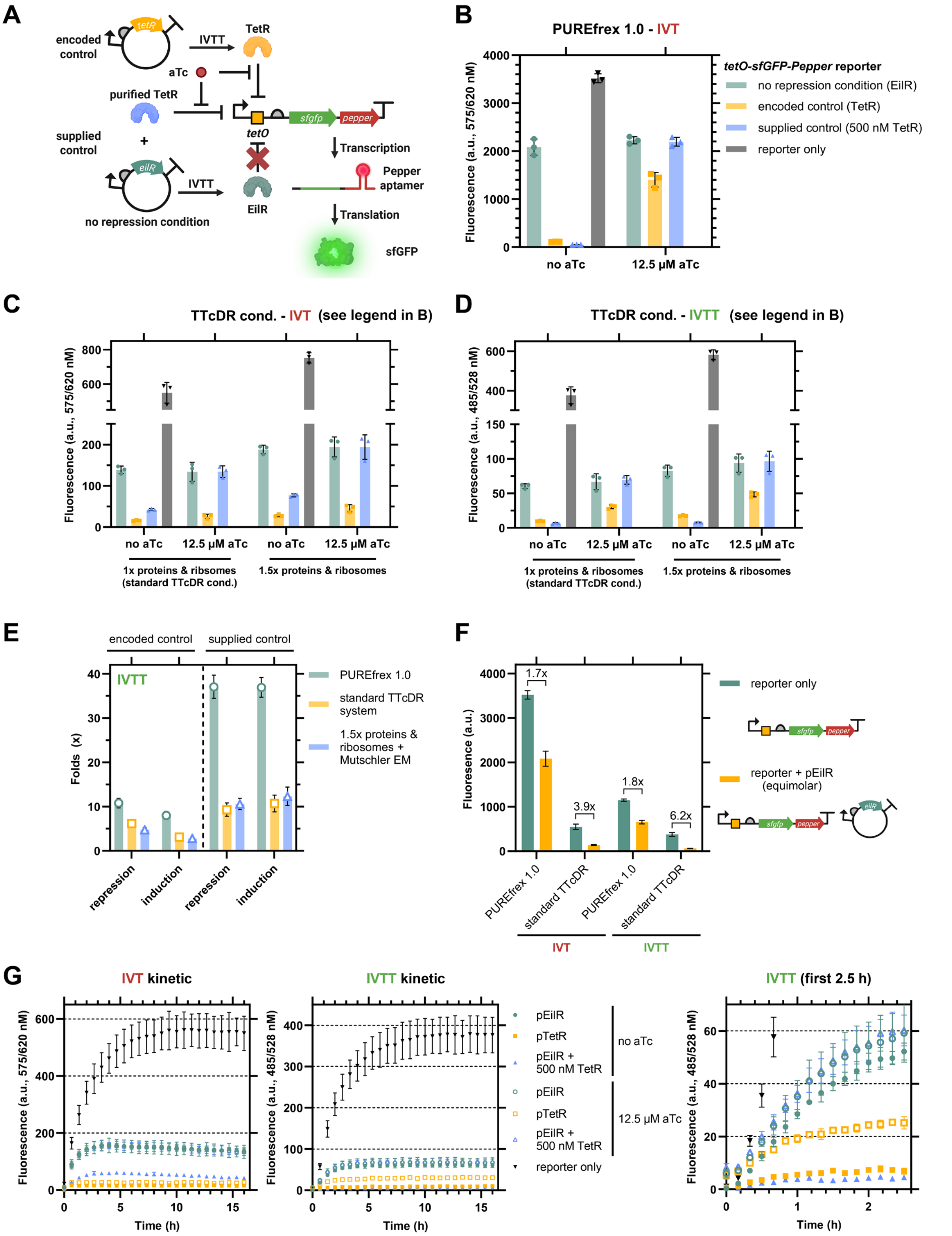

**Figure S5:** Characterization of a genetic circuit with IVTT output. **A:** Schematic of the genetic circuit. TetR is produced from plasmid DNA to repress the $P_{T7}$-*tetO* promoter on the linear DNA template, which controls the expression of the Pepper aptamer and the production of sfGFP. In the presence of anhydrotetracycline (aTc), TetR unbinds $P_{T7}$-*tetO* and the circuit is induced. **B:** 16 h time point data of IVT output (Pepper expression + HBC620 ligand) in PUREfrex 1.0 with either EilR or TetR co-production, or EilR co-production and supplementation of 500 nM purified TetR, or the linear reporter $P_{T7}$-tetO-sfGFP-Pepper alone. **C,D:** Comparison of IVT (**C**) and IVTT (**D**) output (16 h time point) under standard TTcDR conditions and with 1.5x proteins & ribosomes in Mutschler energy mix. **E:** Repression (0 µM aTc) and induction (12.5 µM aTc) folds of the genetic circuit (IVTT output) in PUREfrex 1.0, standard TTcDR conditions and with 1.5x PUREfrex 1.0 proteins and ribosomes + Mutschler energy mix. **F:** Comparison of IVT and IVTT output in PUREfrex 1.0 and under standard TTcDR conditions from the linear reporter template alone and when co-producing EilR from plasmid DNA. The values above the brackets indicate the fold difference between reporter output from the linear DNA template alone and gene co-expression from a circular DNA template, showing that under TTcDR conditions the co-expression of genes from a plasmid DNA reduce the output from the linear template more than in PUREfrex 1.0. **G:** Time course data for the IVT and IVTT outputs of the genetic circuit under standard TTcDR conditions. Data are the mean of n = 3 replicates ± SD.

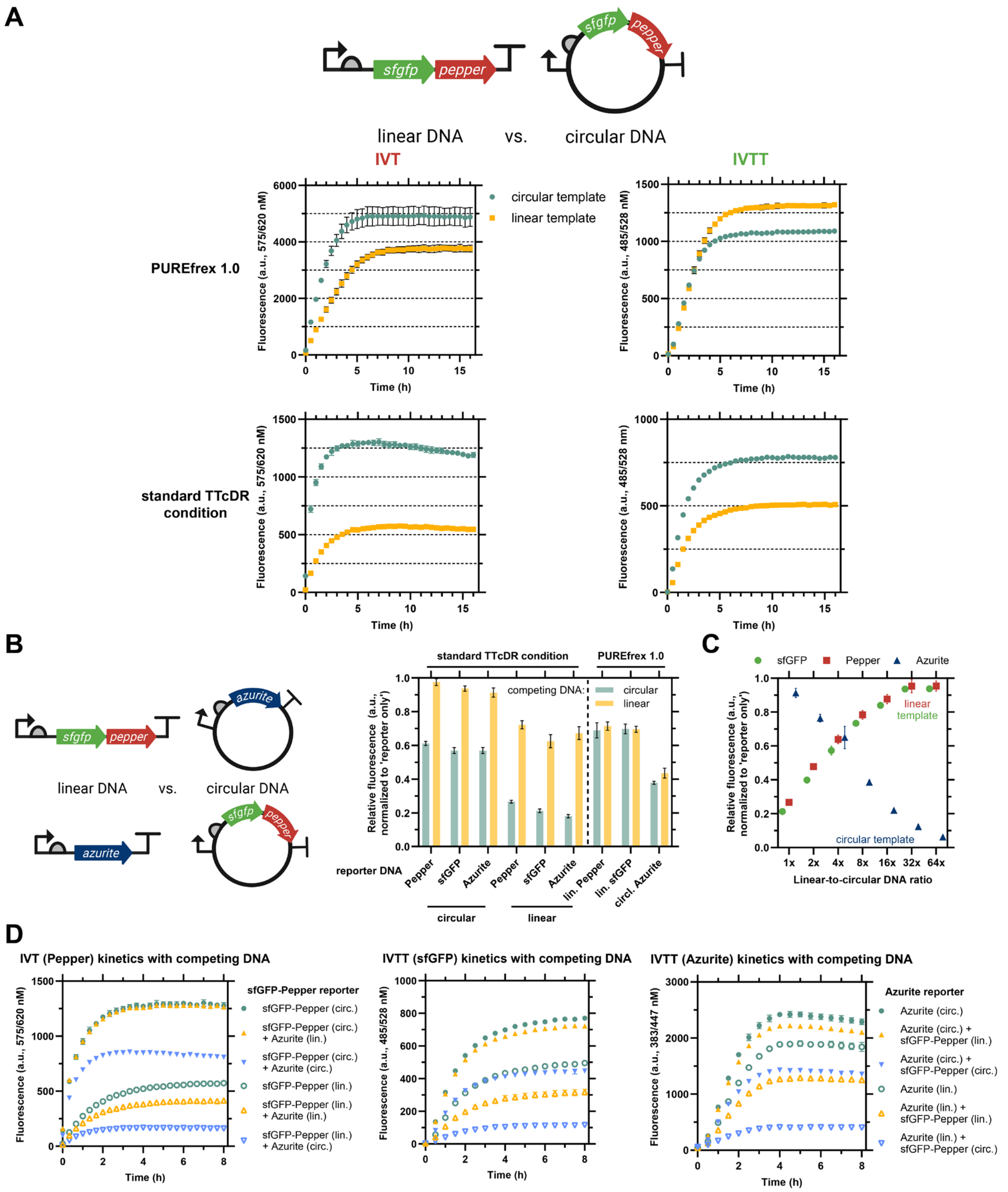

**Figure S6:** Effect of DNA topology on IVT and IVTT output from individual or co-expressing samples. **A:** IVT and IVTT outputs from either linear or circular DNA (see scheme above) in PUREfrex 1.0 and under standard TTcDR conditions. **B:** Competition experiment of a reporter DNA template (either linear or circular, encoding either sfGFP-Pepper or Azurite), and a competing DNA template, either linear or circular, encoding the respective other reporter under standard TTcDR conditions or in PUREfrex 1.0. The fluorescent reporter output in presence of a competing DNA template was normalized to respective reporter only controls. Data are based on 16 h end points. **C**: Titration of linear to circular DNA ratio using a sfGFP-Pepper reporter encoded on linear DNA template and an Azurite reporter encoded on circular DNA template. **D**: Time course data of the competition experiment (in B) of IVT (left), IVTT of sfGFP (center) and IVTT of Azurite (right). Data are the mean of n = 3 replicates ± SD.

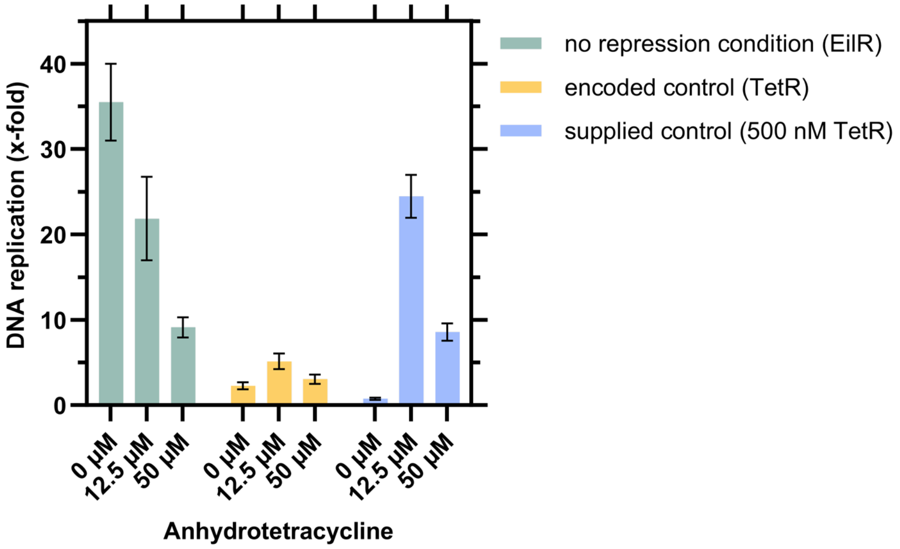

**Figure S7:** Effect of increasing the anhydrotetracycline (aTc) concentration from 12.5 µM to 50 µM. Instead of improving the induction of the genetic circuit, higher concentrations of aTc reduce the DNA replication output of the TTcDR system. Data are from a genetic circuit at 16x linear-to-circular DNA ratio co-producing either TetR (repressing) or EilR (non-repressing) from plasmid DNA (encoded control) with Φ29 DNA polymerase wild-type reporter. For supplied control, 500 nM purified TetR was added to samples co-producing EilR. The genetic circuit was performed either in the absence or in the presence of 12.5 µM or 50 µM aTc. Data are the mean of n = 3 replicates ± SD. TTcDR data are baseline corrected with data from inactive Φ29 DNA polymerase samples (18).

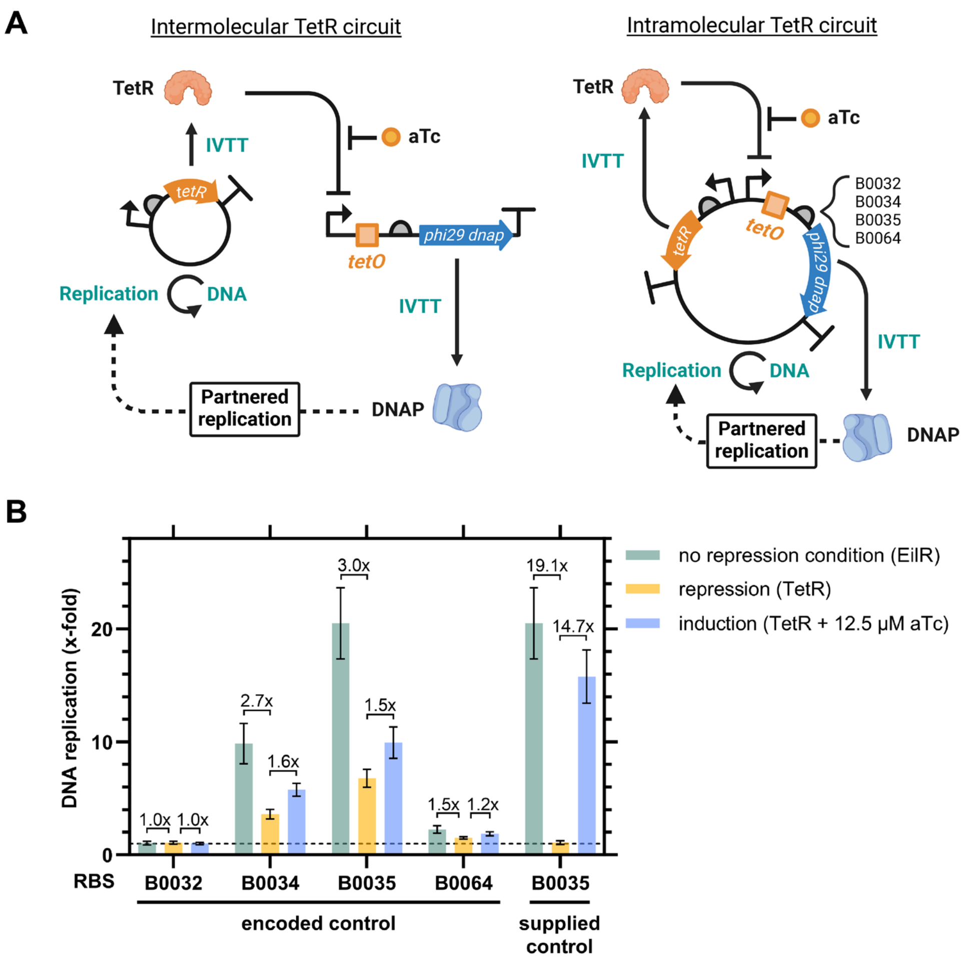

**Figure S8:** Testing an intramolecular TetR circuit with TTcDR output. **A:** Schematic comparison of the intermolecular and intramolecular TetR circuit with TTcDR output. Previously, we tested the genetic circuit using two reporter templates (one linear, one circular). The transcription-translation machinery co-localizes with the respective DNA template molecule (1), so that the produced TetR and phi29 DNA polymerase must diffuse to the other DNA molecule to exert their function. By encoding all genes on one plasmid, we sought to reduce the distance for the proteins to diffuse. See Note S2 for more details on this approach. **B:** TTcDR output with either encoded control (TetR or EilR co-production) or supplemented control (500 nM purified TetR and EilR co-production). The genetic circuit was induced with 12.5 μM anhydrotetracycline (aTc). We tested four ribosome binding site (RBS) sequences that control translation initiation of phi29 DNA polymerase. Fold changes of repression and induction are shown above the bars. Data are the mean of n = 3 replicates ± SD. TTcDR data are baseline corrected with data from inactive Φ29 DNA polymerase samples (18).

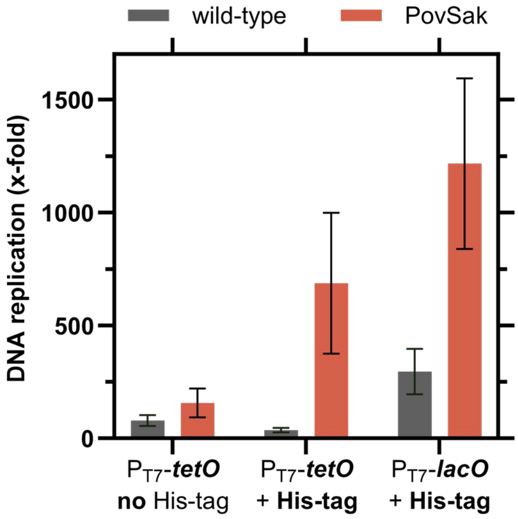

**Figure S9:** Promoter and N-terminal sequence influence DNA replication output. We compared the effect of an N-terminal His-tag on DNA replication by Φ29 DNA polymerase wild-type and the PovSak mutant. In addition, we compared the $P_{T7}$-*tetO* and $P_{T7}$-*lacO* promoters with the His-tagged variants. TTcDR was run with 16x linear-to-circular DNA ratio under standard TTcDR conditions (4 nM linear template encoding the Φ29 DNA polymerase variants and 0.25 nM pGFP-Pepper). Data are the mean of n = 2 replicates ± SD. TTcDR data are baseline corrected with data from inactive Φ29 DNA polymerase samples (18).

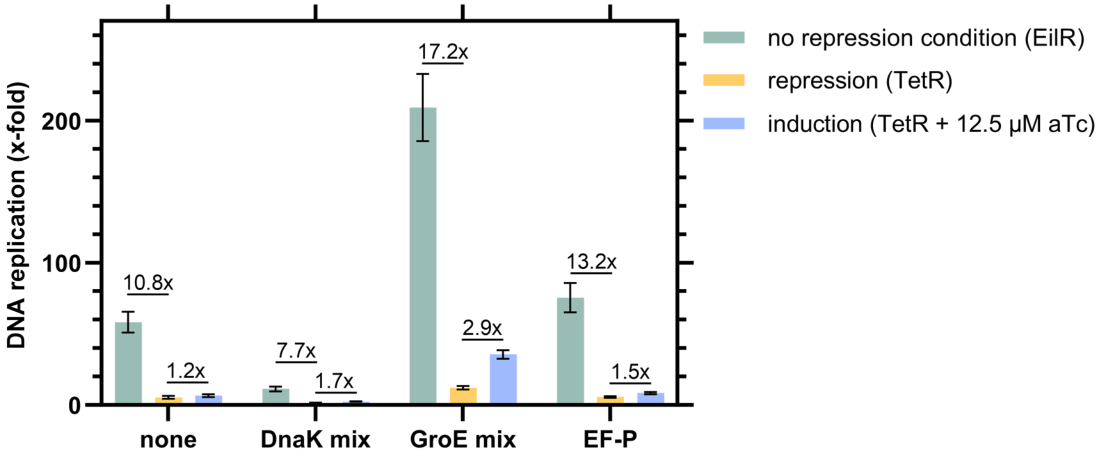

**Figure S10:** Influence of chaperone mixes and EF-P on TetR-based TTcDR circuit performance. TTcDR output with encoded control (TetR or EilR co-production) and wild-type Φ29 DNA polymerase reporter. The genetic circuit was induced with 12.5 µM anhydrotetracycline (aTc). DnaK mix (#PF003), GroE mix (#PF004) and EF-P (#PFS052) have been purchased from GeneFrontier and used according to the vendor's suggestions. Fold changes of repression and induction are shown above the bars. Data are the mean of n = 3 replicates ± SD. TTcDR data are baseline corrected with data from inactive Φ29 DNA polymerase samples (18). See Note S3 for more information.

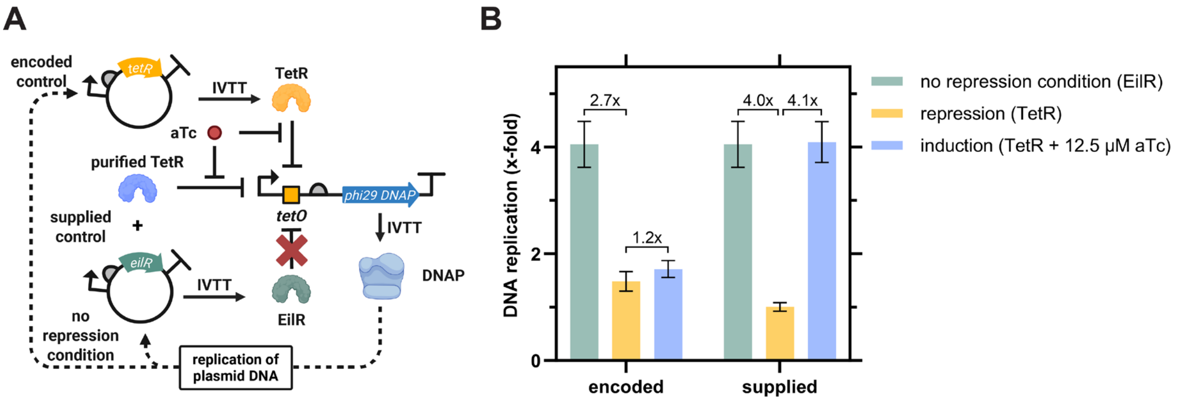

**Figure S11:** Characterization of a TetR-based genetic circuit with TTcDR output at equimolar concentration. **A:** Schematic of the genetic circuit. TetR is produced from plasmid DNA to repress the $P_{T7}$-*tetO* promoter on the linear DNA template, which controls the production of Φ29 DNA polymerase. In the presence of anhydrotetracycline (aTc), TetR unbinds $P_{T7}$-*tetO* and the circuit is induced, leading to the production of DNA polymerase that replicates the plasmid DNA. For encoded control, either TetR (repressing) or EilR (non-repressing) were co-produced. For supplied control, 500 nM purified TetR was added to samples co-producing EilR. **B:** TTcDR output from the genetic circuit at equimolar concentration of 4 nM linear and circular DNA co-producing either EilR (non-repressing) or TetR (repressing). Data are the mean of n = 3 replicates ± SD. TTcDR data are baseline corrected with data from inactive Φ29 DNA polymerase samples (18).

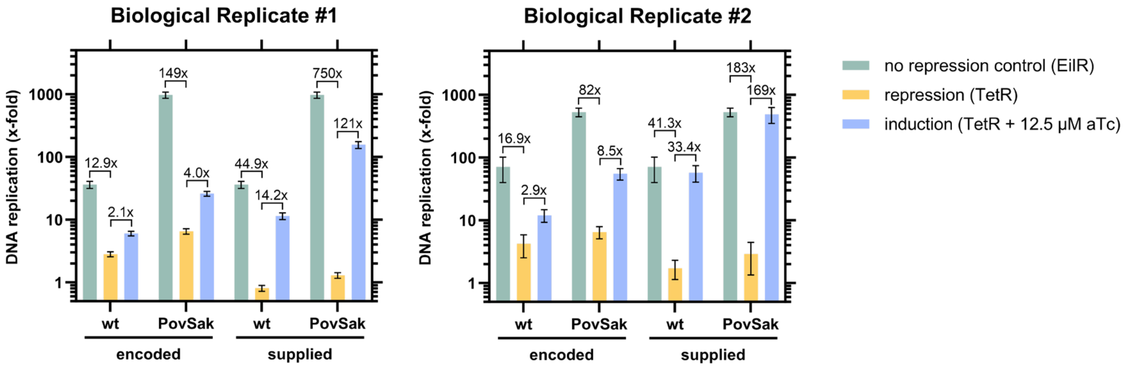

**Figure S12:** Biological replicates of the comparison of Φ29 DNA polymerase wild-type and the His-tagged PovSak mutant in TetR-controlled TTcDR. Biological replicate #1: same as Figure 2F; biological replicate #2 was prepared with all components renewed, i.e. new tRNA, energy mix and DNA preparation, new PUREfrex 1.0 system batch, new batches of each small molecule component, including anhydrotetracycline, and murine RNase inhibitor. Despite overall lower DNA replication in biological replicate #2, the general trends from biological replicate #1 remain. Fold changes of repression and induction are shown above the bars. All samples used 16x linear-to-circular DNA ratios. Data are the mean of n = 3 replicates ± SD. TTcDR data are baseline corrected with data from inactive Φ29 DNA polymerase samples (18).
